# HCR Lateral Flow Assays for Amplified Instrument-Free At-Home SARS-CoV-2 Testing

**DOI:** 10.1101/2022.09.18.508442

**Authors:** Samuel J. Schulte, Jining Huang, Niles A. Pierce

## Abstract

The lateral flow assay format enables rapid, instrument-free, at-home testing for SARS-CoV-2. Due to the absence of signal amplification, this simplicity comes at a cost in sensitivity. Here, we enhance sensitivity by developing an amplified lateral flow assay that incorporates isothermal, enzyme-free signal amplification based on the mechanism of hybridization chain reaction (HCR). The simplicity of the user experience is maintained using a disposable 3-channel lateral flow device to automatically deliver reagents to the test region in three successive stages without user interaction. To perform a test, the user loads the sample, closes the device, and reads the result by eye after 60 minutes. Detecting gamma-irradiated SARS-CoV-2 virions in a mixture of saliva and extraction buffer, the current amplified HCR lateral flow assay achieves a limit of detection of 200 copies/*μ*L using available antibodies to target the SARS-CoV-2 nucleocapsid protein. By comparison, five commercial unamplified lateral flow assays that use proprietary antibodies exhibit limits of detection of 500 copies/*μ*L, 1000 copies/*μ*L, 2000 copies/*μ*L, 2000 copies/*μ*L, and 20,000 copies/*μ*L. By swapping out antibody probes to target different pathogens, amplified HCR lateral flow assays offer a platform for simple, rapid, and sensitive at-home testing for infectious disease. As an alternative to viral protein detection, we further introduce an HCR lateral flow assay for viral RNA detection.

**HCR lateral flow assay:**

- Amplified
- Instrument-free
- At-home
- 60 min
- Naked eye
- SARS-CoV-2
- 200 copies/*μ*L

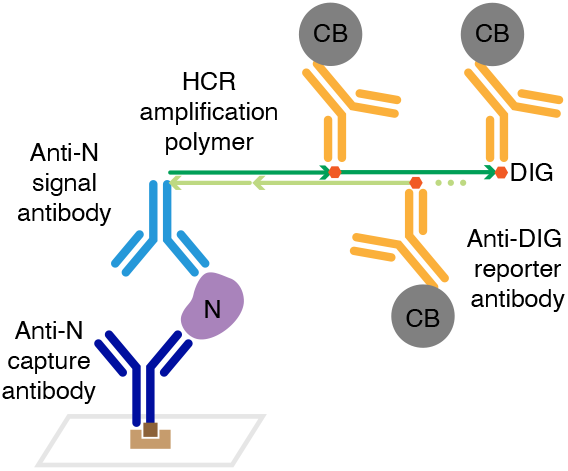

## INTRODUCTION

In March of 2020, the COVID-19 pandemic revealed that lab-based testing could address the technical requirements for detecting the SARS-CoV-2 virus, but could not readily scale to meet the needs of the global population during a pandemic. To address this shortfall in testing capacity, we wondered whether it would be possible to engineer a simple disposable test that could be used at home without special expertise. For inspiration, we looked to disposable at-home pregnancy tests, which had already been in wide use for decades. At-home pregnancy tests employ a lateral flow assay format in which a target protein abundant in urine during early pregnancy moves via capillary forces through a porous substrate, binding in a sandwich between a first antibody carrying a colored label and a second immobilized antibody that concentrates the label within a test region visible to the naked eye.^1^ The resulting signal is unamplified (i.e., one labeled antibody generates signal for one detected target protein), placing limits on sensitivity, but the striking simplicity of lateral flow assays makes them ideal for home use. To take a test, the user simply adds the sample to the disposable device and then checks by eye for a colored signal in the test region after a prescribed number of minutes.

One challenge to developing a lateral flow assay for detection of SARS-CoV-2 virions is their relative scarcity in readily sampled biological fluids. The protein that serves as a pregnancy marker in urine rises to ≈ 10^10^ copies/*μ*L during the first month of pregnancy,^2,3^ with unamplified commercial lateral flow assays typically providing limits of detection of ≈ 10^7^ copies/*μ*L.^3,4^ By comparison, in March 2020, two SARS-CoV-2 studies revealed median viral loads of 158 and 3300 virions/*μ*L in saliva,^5,6^ and lab-based tests using reverse transcription quantitative PCR (PCR tests) achieved limits of detection of 0.1–0.6 copies/*μ*L.^7,8^ Based on these numbers, we set the goal of developing an amplified lateral flow assay that would enable detection of 1000 SARS-CoV-2 virions/*μ*L, representing an increase in sensitivity of approximately four orders of magnitude relative to at-home pregnancy tests. Due to widely reported patient discomfort during nasopharyngeal swabbing for PCR tests, we decided to focus on saliva samples as they are readily obtainable without discomfort or medical expertise.

To boost sensitivity while maintaining simplicity, we hypothesized that signal amplification based on the mechanism of hybridization chain reaction (HCR)^9^ would be well-suited for adaptation to the lateral flow assay format. HCR has been previously used to provide in situ signal amplification for RNA and protein imgaging within fixed biological specimens.^10–13^ In that context, target molecules are detected by probes carrying HCR initiators that trigger chain reactions in which fluorophore-labeled HCR hairpins self-assemble into tethered fluorescent HCR amplification polymers,^10–13^ generating amplified signals in situ at the locations of target molecules within cells, tissue sections, or whole-mount embryos; the specimen is then imaged with a fluorescence microscope to map the expression patterns of target molecules in an anatomical context.^10–15^ HCR signal amplification has critical properties that make it attractive for use in an at-home testing platform: HCR polymerization is isothermal and operates efficiently at room temperature, the resulting amplification polymers are tethered to their initiating probes to concentrate the amplified signal at the target location, and HCR is enzyme-free, employing robust reagents that do not require cold-storage. However, some aspects of HCR imaging protocols presented us with challenges when contemplating at-home use: multiple hands-on steps (probe addition, probe incubation, and probe removal via washing, followed by amplifier addition, amplifier incubation, and amplifier removal via washing), protocol duration (typically overnight probe incubation and overnight amplifier incubation), and the need for a fluorescence microscope to image the results. To eliminate the need for hands-on steps, we planned to attempt the use of multi-channel lateral flow devices to automatically deliver reagents to the test region in successive stages.^16,17^ To dramatically speed up signal amplification, we planned to work at higher reagent concentrations than are typical for HCR imaging experiments. And to eliminate the need for a fluorescence microscope, we planned to switch to colored rather than fluorescent reporters, which are bulky by comparison (potentially even larger than the HCR hairpins themselves), but can be seen by the human eye if concentrated in the test region in sufficient abundance.

As a precursor reality check, we verified that by increasing the HCR hairpin concentration, HCR amplification polymers can grow to a length of over 500 hairpins within 10 minutes (Figure S18), matching the two orders of magnitude of signal amplification achieved in situ using overnight amplification for HCR imaging.^11,13^ We then set out to pursue two parallel projects developing amplified HCR lateral flow assays for detecting either viral protein or viral genomic RNA, uncertain which approach would be more effective.

Seeking to maintain the attractive properties of existing pregnancy tests while addressing the more demanding challenge of SARS-CoV-2 detection, we set firm design criteria:

- *Simple*: from the user’s perspective, the test should be as simple to use as a pregnancy test, enabling routine at-home use by a non-expert.
- *Inexpensive*: the test device should be disposable and not require at-home instrumentation.
- *Robust*: the test should avoid reagents (e.g., enzymes) that require cold storage.
- *Rapid*: the test should return results in 1 hour or less.
- *Sensitive*: the test should have a limit of detection of 1000 virions/*μ*L or lower.

During the two years that we have been working to achieve these goals, the testing landscape has evolved. While lab-based PCR tests remain the gold standard for SARS-CoV-2 testing, a number of unamplified lateral flow assays have been commercialized for at-home testing. These tests are simple, inexpensive, robust, and rapid, and are highly reliable when they return a positive result (e.g., 96–100%),^18,19^ but sensitivity limitations can lead to a high false-negative rate (e.g., 25–50% in two hospital studies).^18,19^ With this work we seek to partially bridge the sensitivity gap between commercial unamplified lateral flow assays and lab-based PCR tests so as to reduce the false-negative rate for at-home testing. Other efforts to enhance sensitivity by introducing signal amplification into SARS-CoV-2 lateral flow assays, including use of loop-mediated isothermal amplification (LAMP)^20,21^ and CRISPR/Cas,^22,23^ have led to compromises on simplicity, cost, and/or robustness, requiring multiple user steps, dedicated instrumentation, and/or enzymes with strict storage requirements.

Here, we have developed an amplified HCR lateral flow assay for SARS-CoV-2 protein detection that is simple, disposable, enzyme-free, returns a result in 1 hour, and achieves a limit of detection lower than all five commercial SARS-CoV-2 rapid antigen tests that we evaluated. On the other hand, in developing an amplified HCR lateral flow assay detecting the SARS-CoV-2 RNA genome, we have so far found it necessary to include a heat extraction step, increasing the assay complexity and time, while matching the enhanced sensitivity of our protein test. Because the RNA test employs DNA signal probes that can be quickly redesigned to target a new pathogen in the case of an emerging infectious disease, it offers a novel approach to at-home testing that merits further study and refinement. In the case of protein detection, amplified HCR lateral flow assays fill an important sensitivity gap between current commercial lateral flow assays and PCR tests.

## RESULTS AND DISCUSSION

### Viral Protein Detection

To detect viral protein, we target the same SARS-CoV-2 nucleocapsid protein (N) that is targeted by numerous commercial SARS-CoV-2 lateral flow assays. The protein target N decorates the RNA genome within the viral envelope with ≈ 10^3^ copies/virion^24^ (enhancing sensitivity) and is strongly immunogenic^25,26^ (facilitating development of high-affinity anti-N antibodies). In a conventional lateral flow assay (Figure 1a), the target protein is detected in a sandwich between a reporter-labeled signal antibody that binds a first target epitope and a capture antibody that binds a second target epitope, immobilizing the signal in the test region when the target is present in the sample.^1^ If the target is sufficiently abundant, the signal in the test region is visible to the naked eye. To incorporate HCR into an amplified lateral flow assay for SARS-CoV-2 (Figure 1b), the anti-N signal antibody is instead labeled with one or more HCR initiators. After the antibody/target sandwich is immobilized in the test region, the HCR initiators labeling the anti-N signal antibody trigger the self-assembly of HCR hairpins into tethered HCR amplification polymers. For fluorescence imaging applications, HCR hairpins are fluorophore-labeled for imaging with a fluorescence microscope,^10–13^ but for lateral flow assays, a colored label is required to enable detection by the human eye. In order to maximize the signal per HCR hairpin while avoiding impeding polymerization kinetics by labeling hairpins with bulky colored reporters, we instead label HCR hairpins with a hapten (digoxigenin; DIG), which in turn is detected by an anti-DIG reporter antibody carrying carbon black (CB). The anti-N capture antibody is biotinylated and is itself captured in the test region by pre-immobilized polystreptavidin R (PR).^27,28^

**Figure 1.**
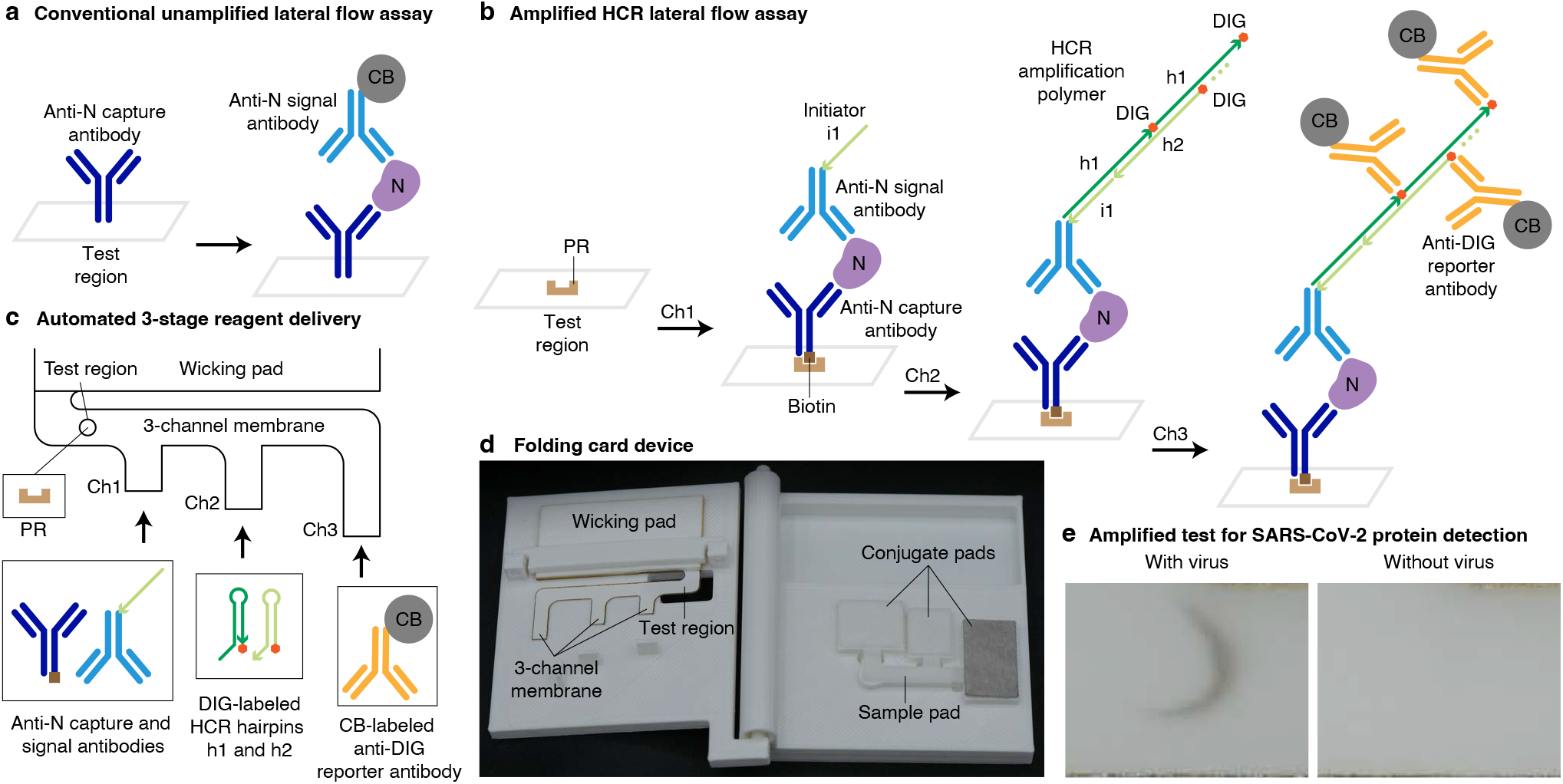
Amplified HCR lateral flow assay for SARS-CoV-2 via detection of nucleocapsid protein (N). (a) Conventional unamplified lateral flow assay. Each target protein N is immobilized in the test region in a sandwich between an anti-N capture antibody and a CB-labeled anti-N signal antibody, generating one unit of signal per target. N: nucleocapsid protein. CB: carbon black. (b) Amplified HCR lateral flow assay. Each target protein N is immobilized by PR in the test region in a sandwich between a biotinylated anti-N capture antibody and an anti-N signal antibody labeled with HCR initiator i1, triggering self-assembly of DIG-labeled HCR hairpins (h1 and h2) to form a tethered HCR amplification polymer decorated with DIG that is subsequently bound by multiple CB-labeled anti-DIG reporter antibodies, generating multiple units of signal per target. PR: polystreptavidin R. DIG: digoxigenin. (c) Automated delivery of reagents to the test region from Channels 1, 2, and 3 in succession using a 3-channel membrane. (d) Folding card device. The left page of the device contains the 3-channel membrane and wicking pad. The right page of the device contains three conjugate pads (containing dried reagents for Channels 1, 2, and 3) and a sample pad. To perform the test, the user adds the sample to the sample pad, closes the device, and reads the result after 60 min. (e) SARS-CoV-2 test with or without gamma-irradiated virus spiked into a mixture of saliva and extraction buffer at 1000 copies/*μ*L.

For conventional unamplified lateral flow assays, the protein target and anti-target signal antibodies flow to the test region along a single membrane channel. For our amplified HCR lateral flow assay, we leverage prior work that explored the use of multi-channel lateral flow assays.^16,17^ Reagents are automatically delivered to the test region in three successive stages using a 3-channel membrane in which channels of different lengths lead into a unified channel before reaching the test region (Figure 1c). The anti-N signal and capture antibodies bind the target protein N and travel via the shortest membrane channel (Channel 1) to reach the test region first, where the antibody/target sandwich is immobilized via binding of biotinylated anti-N capture antibodies to pre-immobilized PR. DIG-labeled HCR hairpins travel via a channel of intermediate length (Channel 2) and reach the test region next, where initiators on the anti-N signal antibodies trigger growth of tethered DIG-labeled HCR amplification polymers. CB-labeled anti-DIG reporter antibodies travel via the longest channel (Channel 3) and arrive in the test region last, where they decorate the HCR amplification polymers to generate an amplified colored signal in the test region. Reagents for each channel are dried onto separate conjugate pads, which are rehydrated simultaneously when the user adds the sample to the sample pad. Upon rehydration, successive delivery of the reagents to the test region occurs automatically without user interaction, as draining of the first conjugate pad frees the unified channel for draining of the second conjugate pad, which in turn frees the unified channel for draining of the third conjugate pad.^16^ Our prototype device takes the form of a folding card (Figure 1d). The right page of the card contains the sample pad and three conjugate pads. The left page of the card contains the 3-channel membrane and the wicking pad, which absorbs liquid to induce continued capillary flow through the channels. The left page is functionalized with three prongs which disconnect the sample pad from the conjugate pads upon folding the card, limiting the volume that flows from each conjugate pad and preventing flow between conjugate pads.

To perform a test, the user adds saliva to a tube containing extraction buffer (disrupting the viral envelope to expose the protein targets N), adds the extracted sample to the sample pad, and closes the card to create contact between the membrane and the three conjugate pads, initiating the consecutive flow of liquid from Channels 1,2, and 3 (Supplementary Movie 1). After 60 minutes, the user reads either a positive result (black signal) or a negative result (no signal) in the test region with the naked eye (Figure 1e). Due to automated multi-channel reagent delivery, this amplified HCR lateral flow assay retains the simplicity of conventional commercial unamplified lateral flow assays, requiring only sample addition and card closure before reading the result, with signal amplification occurring unbeknownst to the user.

To characterize sensitivity, we ran HCR lateral flow assays on gamma-irradiated SARS-CoV-2 virus spiked into a mixture of saliva and extraction buffer at a range of concentrations, revealing a limit of detection of 200 virus copies/*μ*L (Figure 2a). No background staining was observed in the test region for experiments run without spiked-in virus (Figure 2b). To characterize cross-reactivity, experiments were run with spiked-in recombinant N protein from a different betacoronavirus (OC43) or with spiked-in nucleoprotein from Influenza Type A (H3N2); no staining was observed in the test region in either case even with off-target proteins at high concentration (equivalent to ≈ 10^6^ virions/*μ*L;^24,29^ Figure 2c).

**Figure 2.**
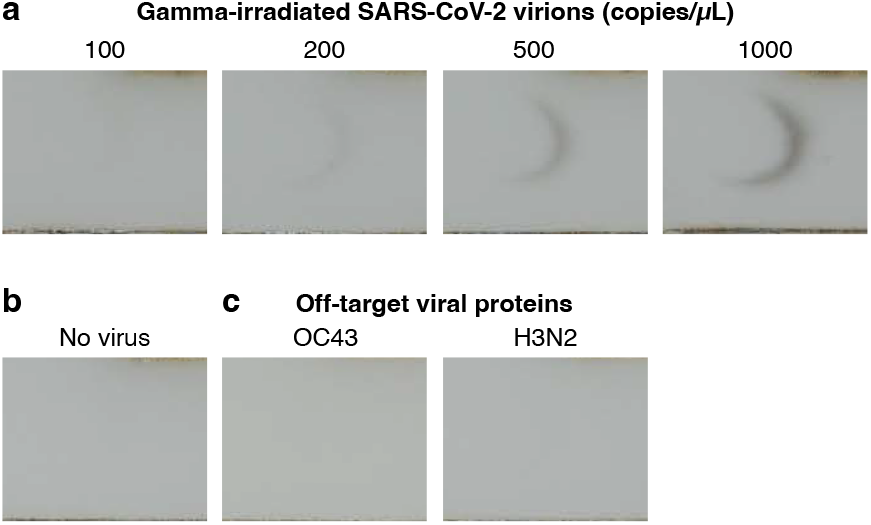
Amplified HCR lateral flow assay performance for SARS-CoV-2 via detection of nucleocapsid protein (N). (a) Sensitivity: gamma-irradiated SARS-CoV-2 spiked into a mixture of saliva and extraction buffer at different concentrations revealed a limit of detection of 200 copies/*μ*L. (b) Background: no staining observed in the absence of virus. (c) Cross-reactivity: no staining observed for N protein from a different betacoronavirus (OC43; 83.74 ng/mL) or nucleoprotein from Influenza Type A (H3N2; 50.43 ng/mL) spiked into a mixture of saliva and extraction buffer. See Figures S9 and S10 for replicates.

Assay performance was then benchmarked against five commercial SARS-CoV-2 lateral flow assays by spiking gamma-irradiated SARS-CoV-2 into the extraction buffer included with each test kit and loading the extracted sample according to the manufacturer’s instructions. Each test was performed in triplicate at each concentration, and the limit of detection for a given kit was defined as the lowest tested concentration for which all three replicates had a visible signal. While the amplified HCR lateral flow assay detects gamma-irradiated SARS-CoV-2 at 200 copies/*μ*L, none of the unamplified commercial tests were able to detect SARS-CoV-2 at this concentration. The limits of detection for the five kits were 500 copies/*μ*L, 1000 copies/*μ*L, 2000 copies/*μ*L, 2000 copies/*μ*L, and 20,000 copies/*μ*L (see Table 1 for a summary and Figures S9 and S11–S15 for replicate images). It is important to note that these commercial tests use proprietary anti-N antibodies, the affinity of which play a critical role in assay sensitivity.^27^ Nevertheless, despite not having access to proprietary antibodies utilized by the test kit manufacturers, the HCR-amplified assay still achieves a lower limit of detection for SARS-CoV-2.

**Table 1.** Test results for SARS-CoV-2 rapid tests detecting nucleocapsid protein (N): amplified HCR lateral flow assay vs five commercial unamplified lateral flow assays. Gamma-irradiated virus spiked into a mixture of saliva and extraction buffer (current work) or manufacturer-provided extraction buffer (commercial tests). *N* = 3 replicates for each concentration. Each replicate was judged by eye for a positive 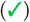 or negative 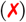 test result. Not tested (n.t.). See Figures S9 and S11–S15 for images.

| Test | Gamma-irradiated virus copies/μL | | | | | | | | Anti-N antibodies | Amplification | Duration (min) |
| --- | --- | --- | --- | --- | --- | --- | --- | --- | --- | --- | --- |
|  | 100 | 200 | 500 | 1,000 | 2,000 | 5,000 | 10,000 | 20,000 |  |  |  |
| Current work | XXX | ✓✓✓ | ✓✓✓ | ✓✓✓ | n.t. | n.t. | n.t. | n.t. | Available | HCR | 60 |
| Commercial A | XXX | XXX | XXX | XXX | ✓✓✓ | n.t. | n.t. | n.t. | Proprietary | None | 15 |
| Commercial B | XXX | XXX | XXX | XXX | ✓✓✓ | n.t. | n.t. | n.t. | Proprietary | None | 10 |
| Commercial C | XXX | XXX | ✓✓✓ | ✓✓✓ | n.t. | n.t. | n.t. | n.t. | Proprietary | None | 15 |
| Commercial D | XXX | XXX | XXX | XXX | XXX | XXX | XXX | ✓✓✓ | Proprietary | None | 15 |
| Commercial E | XXX | XXX | XXX | ✓✓✓ | ✓✓✓ | n.t. | n.t. | n.t. | Proprietary | None | 10 |

To quantify the amplification gain provided by HCR in a lateral flow context, we compared the signal using both HCR hairpins (h1 and h2) to the signal using only hairpin h1. In the h1-only condition, polymerization cannot proceed beyond the binding of h1 to the initiator, emulating the unamplified signal of a conventional lateral flow assay where each detected target generates one detectable signal. The amplification gain, calculated as the ratio of amplified to unamplified signal intensities, is 13.7 ± 0.8 (mean ± estimated standard error of the mean via uncertainty propagation for *N* = 3 replicate assays for each experiment type; Figure S20 and Table S2).

This amplified HCR lateral flow assay fulfills the five design requirements that we set in March 2020. Despite the incorporation of signal amplification into the assay, the test remains simple to use. Signal amplification occurs automatically using a 3-channel design, increasing sensitivity while remaining as simple as conventional unamplified lateral flow assays from the user’s perspective. The inexpensive card device and reagents (PR, initiator-labeled anti-N signal antibody, biotinylated anti-N capture antibody, DIG-labeled HCR hairpins, and CB-labeled reporter antibody) are disposable, comparable in cost to those for commercial lateral flow assays, and require no dedicated instrumentation beyond the human eye for readout. The reagents are robust and do not have cold-storage requirements. Despite the extra time required for successive automated delivery of reagents in three stages, the test remains rapid, delivering a result in 60 minutes. Finally, the assay is sensitive, enabling detection of 200 copies/*μ*L of gamma-irradiated SARS-CoV-2 virus. This limit of detection is 2.5× to 100× lower than the limits of detection of 5 commercial SARS-CoV-2 lateral flow assays despite our lack of access to proprietary antibodies.

### Viral RNA Detection

To detect viral RNA, we target the same SARS-CoV-2 single-stranded RNA genome that is detected by lab-based PCR tests (Figure 3a). The target RNA is detected by DNA signal probes complementary to different subsequences along the ≈30,000 nt target, avoiding subsequences shared by other coronaviruses (with the exception of SARS-CoV, which has high sequence similarity to SARS-CoV-2, causes severe disease, and is not circulating^30^). To automatically suppress background that could otherwise arise from non-specific probe binding, our DNA signal probes take the form of split-initiator probe pairs with an HCR initiator split between a pair of probes.^12^ As a result, any individual probe that binds non-specifically will not trigger HCR, but specific hybridization of a pair of probes to adjacent cognate binding sites along the target RNA will colocalize a full HCR initiator capable of triggering HCR signal amplification. To maximize sensitivity, the signal probe set comprises 198 splitinitiator DNA probe pairs.

**Figure 3.**
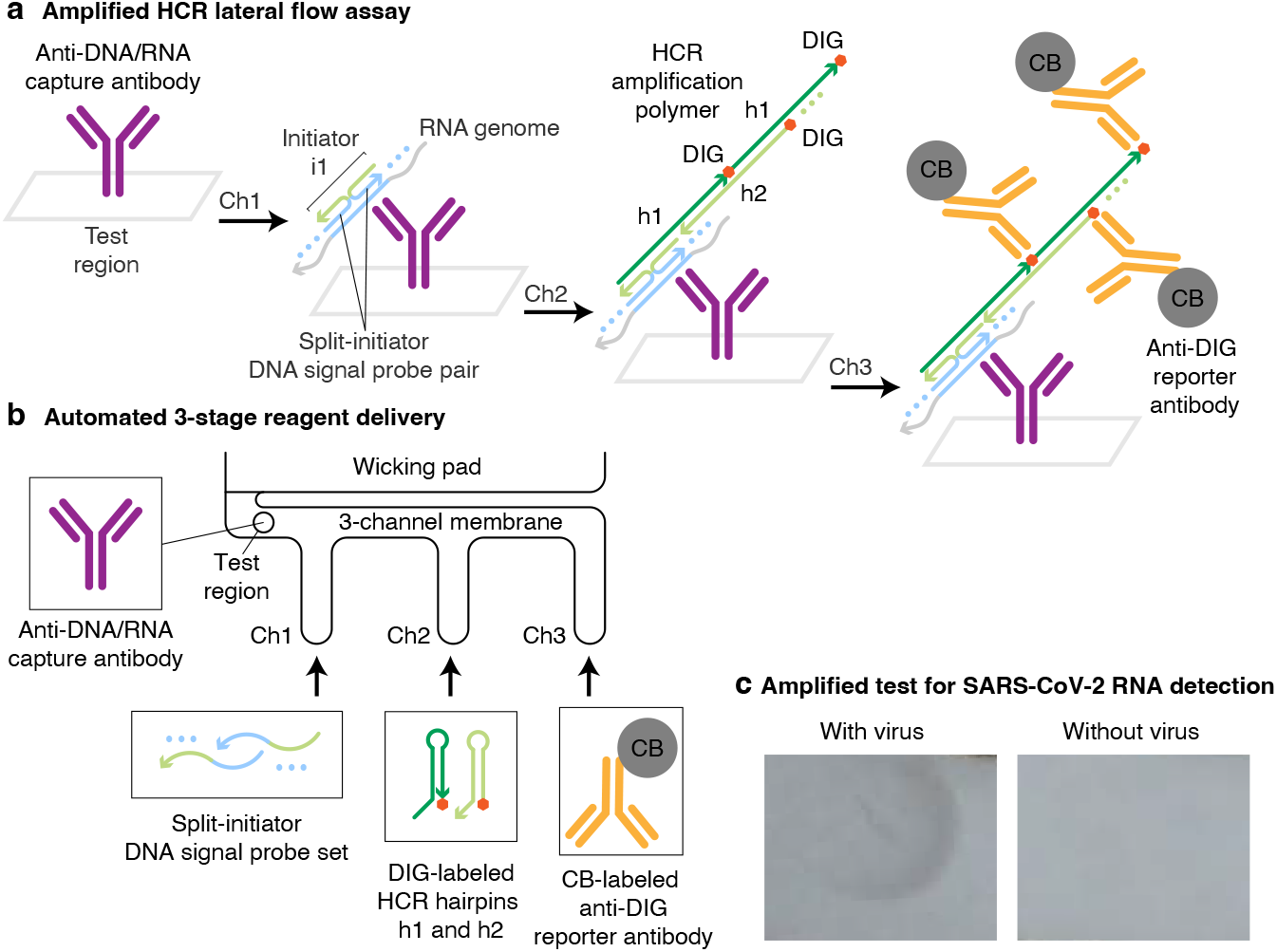
Amplified HCR lateral flow assay for SARS-CoV-2 via detection of the viral RNA genome. (a) Amplified HCR lateral flow assay. Split-initiator DNA signal probe pairs hybridize to cognate binding sites on the viral RNA genome to colocalize full initiator i1; the resulting DNA/RNA duplex is captured in the test region by anti-DNA/RNA capture antibodies; initiator i1 triggers self-assembly of DIG-labeled HCR hairpins (h1 and h2) to form a tethered HCR amplification polymer that is subsequently bound by multiple CB-labeled anti-DIG reporter antibodies, generating multiple units of signal per target. DIG: digoxigenin. CB: carbon black. (b) Automated delivery of reagents to the test region from Channels 1, 2, and 3 in succession using a 3-channel membrane. (c) SARS-CoV-2 test with or without gamma-irradiated virus spiked into extraction buffer at 1000 copies/*μ*L.

By hybridizing to the RNA target, the DNA signal probes create a DNA/RNA duplex at each probe binding site. The probe-decorated target is captured in the test region by an immobilized anti-DNA/RNA capture antibody^31^ that binds to DNA/RNA duplexes. Note that while the capture antibody binds DNA/RNA duplexes independent of sequence, any captured RNA that does not include specifically bound split-initiator DNA signal probe pairs will not trigger HCR, and hence will not contribute to background. After immobilization of the probe-decorated target in the test region, colocalized full HCR initiators trigger the self-assembly of DIG-labeled HCR hairpins into HCR amplification polymers decorated with DIG, which are in turn bound by CB-labeled anti-DIG reporter antibodies. As for our viral protein HCR lateral flow assay, reagents are automatically delivered to the test region in three successive stages using a 3-channel membrane (Figure 3b). The DNA signal probes bind the RNA target and travel via the shortest membrane channel (Channel 1) to reach the test region first, where the probe-decorated target is captured by pre-immobilized anti-DNA/RNA capture antibodies. DIG-labeled HCR hairpins travel via a channel of intermediate length (Channel 2) and reach the test region next, where colocalized full HCR initiators formed by specifically bound split-initiator signal probe pairs trigger growth of tethered DIG-labeled HCR amplification polymers. CB-labeled anti-DIG reporter antibodies travel via the longest channel (Channel 3) and arrive in the test region last, where they bind the HCR amplification polymers to generate an amplified colored signal in the test region.

Extraction of the viral RNA genome requires mild denaturing conditions to remove the viral envelope and the nucleocapsid (N) proteins decorating the RNA genome. These denaturing conditions also help to disrupt native secondary structure in the single-stranded RNA genome prior to detection by the DNA signal probe set. We encountered difficulties incorporating chemical denaturants into the rapid test platform, as the same denaturants that enable target extraction and denaturation also destabilize the DNA/RNA duplexes that underly target detection. By contrast, heat denaturation can be applied to the sample transiently and then removed to allow DNA probe binding.^31^ To date, we have found incorporation of a heat denaturation step to be essential for detection of the SARS-CoV-2 RNA genome in the context of an amplified HCR lateral flow assay. Unfortunately, the addition of a heating step means that we are not yet able to meet our goal that the viral RNA test be as simple to perform as a pregnancy test.

Nonetheless, conceding for the time being that a heat denaturation step is required for the viral RNA lateral flow assay, here we demonstrate a prototype approach using reagents in solution rather than dried onto conjugate pads (i.e., using a half-strip assay format) to facilitate heating the sample/probe mixture on a heat block prior to starting the lateral flow assay. For consistency, the DIG-labeled HCR hairpins and CB-labeled anti-DIG reporter antibodies are also prepared in solution, though these components could alternatively be dried onto conjugate pads. To perform a test, the sample is added to extraction buffer containing the DNA signal probes (Channel 1 reagents), heated to 65 °C for 15 minutes, and then loaded into a well (Channel 1 well) on a 96-well plate proximal to wells containing the DIG-labeled HCR hairpins (Channel 2 well) and CB-labeled anti-DIG reporter antibodies (Channel 3 well; Figure 3b). To start the lateral flow assay, the ends of the three membrane channels are simultaneously submerged into the three wells, leading to automated successive delivery of the Channel 1, 2, and 3 reagents to the test region without user interaction. After 90 minutes, the result is read as either a positive result (black signal) or a negative result (no signal) in the test region with the naked eye (Figure 3c). The 90-minute duration of this lateral flow assay is longer than the 60-minute duration for the protein detection case both because the spacing between the wells on a 96-well plate dictated longer channels, and also because our use of larger reagent volumes delayed the transitions between channel flows (Supplementary Movie 2).

To characterize sensitivity, gamma-irradiated SARS-CoV-2 virus was spiked into extraction buffer with DNA signal probes, heated to 65 °C for 15 min, and run on the amplified HCR lateral flow assay, revealing a limit of detection of 200 virions/*μ*L (Figure 4a). No background staining was observed in the test region for experiments run without spiked-in virus (Figure 4b). To characterize cross-reactivity, experiments were run with synthetic RNA genomes from other coronaviruses spiked in at high concentration (7,200 copies/*μ*L for 229E and 10,000 copies/*μ*L for HKU1); no staining was observed in the test region in either case (Figure 4c). By matching the limit of detection of our viral protein test, this viral RNA test is likewise more sensitive than all five commercial lateral flow assays that we evaluated for viral protein detection, with the caveat that this RNA test uses a half-strip assay format (with reagents in solution rather then dried onto conjugate pads). For the RNA detection setting, the HCR amplification gain was 10 ± 3 (mean ± estimated standard error of the mean via uncertainty propagation for *N* = 3 replicate assays for each experiment type; Figure S21 and Table S2), measured by comparing the signal intensity for assays run using both HCR hairpins (h1 and h2) to assays run using only hairpin h1.

**Figure 4.**
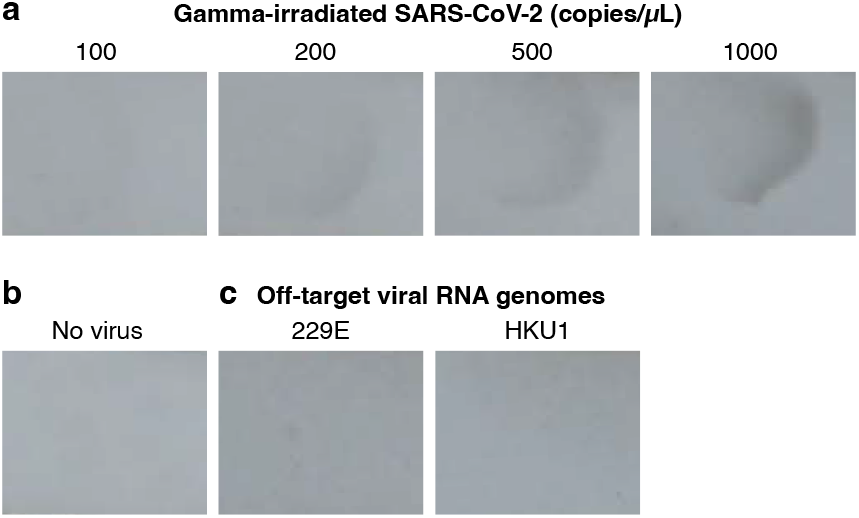
Amplified HCR lateral flow assay performance for SARS-CoV-2 via detection of the RNA genome. (a) Sensitivity: gamma-irradiated SARS-CoV-2 spiked into extraction buffer at different concentrations revealed a limit of detection of 200 copies/*μ*L. (b) Background: no staining observed in the absence of virus. (c) Cross-reactivity: no staining observed for extraction buffer spiked with synthetic RNA genomes from different coronaviruses 229E (7,200 copies/*μ*L) or HKU1 (10,000 copies/*μ*L). See Figures S16 and S17 for replicates.

RNA detection with on-strip HCR amplification introduces a new approach to lateral flow testing for infectious disease, as to our knowledge, all current unamplified commercial lateral flow assays detect protein rather than nucleic acids. An important benefit of RNA detection is the ability to design and synthesize a new DNA signal probe set within a matter of days following sequencing of a new RNA target of interest. As a consequence, shortly after the emergence of a new pathogen, an amplified HCR lateral flow assay could be developed and tested. This capability would fill a critical gap in the at-home testing infrastructure while antibodies suitable for a protein-detection lateral flow assay are developed and screened.

Despite not fulfilling our five design criteria, viral RNA detection with an amplified HCR lateral flow assay demonstrates promise. Currently, the addition of a heating step undermines our goals in several regards, reducing simplicity by adding an extra step, requiring the use of a heat block (which can be viewed as a primitive form of instrumentation), and increasing the test duration. Further testing of chemical denaturants, as well as strategies for automated removal of denaturants, may eliminate the need for a heat denaturation step, enhancing the assay in multiple regards. Nonetheless, with the exception of the heating step, the assay remains simple, exploiting a 3-channel membrane to automatically perform HCR signal amplification without user interaction. Sensitivity is enhanced compared to unamplified commercial SARS-CoV-2 viral protein tests, achieving a limit of detection for gamma-irradiated SARS-CoV-2 of 200 copies/*μ*L. The reagents (anti-DNA/RNA capture antibody, DNA signal probes, DIG-labeled HCR hairpins, CB-labeled anti-DIG reporter antibodies) are inexpensive, and only the human eye is needed to read out the result. If the heating step can be removed, it appears feasible to meet all five design requirements using a full-strip lateral flow assay format (with reagents dried onto conjugate pads as for the protein-detection assay). Even if the heating step cannot be removed, RNA-detection HCR lateral flow assays may still prove useful due to their suitability for rapid deployment of new tests in the face of emerging pathogens.

### Conclusion

Routine at-home testing with an amplified lateral flow assay could be transformative in preventing infectious disease transmission during a pandemic. For example, data and modeling suggest that more than half of SARS-CoV-2 infections are spread unknowingly by asymptomatic carriers.^32, 33^ To enable routine at-home testing for SARS-CoV-2, we have developed an amplified HCR lateral flow assay for viral protein detection that is simple to use (comparable to a pregnancy test), inexpensive (using a disposable device with readout via the naked eye), robust (enzyme-free, using no reagents that require cold-storage), rapid (delivering a result in 60 minutes), and sensitive (detecting 200 copies/*μ*L of gamma-irradiated SARS-CoV-2 in a mixture of saliva and extraction buffer). By comparison, five unamplified commercial lateral flow assays exhibited limits of detection that are 2.5×, 5×, 10×, 10×, and 100× higher than our amplified HCR lateral flow assay, despite the advantage of using proprietary antibodies. Current commercial unamplified SARS-CoV-2 lateral flow tests lead to high false-negative rates,^18,19^ indicating that their limits of detection fall not in the lower tail of the distribution of clinical viral loads, but near the middle of the distribution where further reductions of the limit of detection will be maximally impactful in reducing the false-negative rate. While we focused our assay development on saliva due to its convenience relative to nasopharyngeal swabbing, anterior nasal swabbing is now used for numerous commercial SARS-CoV-2 tests and offers another convenient alternative.

This work demonstrates that it is possible to combine the enhanced sensitivity of HCR signal amplification with the simplicity of the lateral flow assay format, which was achieved using a 3-channel membrane to automatically deliver reagents to the test region in three successive stages without user interaction. In the future, it will be desirable to use best-in-class antibodies so that the enhanced sensitivity of HCR signal amplification pushes the limit of detection of the HCR lateral flow assay even closer to that of PCR tests. With further optimization of HCR in the context of automated lateral flow reagent delivery, there is also the potential to increase the HCR signal gain from the current one order of magnitude to the two orders of magnitude achieved in HCR imaging applications. While the 60-minute run time of our amplified test is higher than the 10-or-15-minute run time of unamplified commercial lateral flow assays, we anticipate that in many situations, users will prefer a test that offers superior sensitivity while still providing a result in 1 hour. By switching out SARS-CoV-2 antibodies for antibodies targeting other pathogens, amplified HCR lateral flow assays offer a versatile platform for sensitive at-home testing, including for emerging pathogens. Amplified HCR lateral flow assays for viral RNA detection have not yet achieved our goals for simplicity and rapidity, but represent a promising new approach for infectious disease testing and would enable nimble assay development upon the sequencing of novel pathogens.

## METHODS SUMMARY

### Viral Protein Detection

The disposable folding card device was printed with a 3D printer. The nitrocellulose membrane and wicking pad were overlapped on an adherent backing material and cut into a 3-channel geometry with a laser cutter. The sample pad was cut with a laser cutter, blocked, dried, and adhered to the right page of the card device. The conjugate pads were cut with a laser cutter, blocked, loaded with reagents (Channel 1: anti-N signal and capture antibodies; Channel 2: DIG-labeled HCR hairpins h1 and h2; Channel 3: CB-labeled anti-DIG reporter antibody), dried, and adhered to the right page of the device. The membrane was spotted with PR in the test region, dried, and adhered to the left page of the device. To run the assay, gamma-irradiated SARS-CoV-2 virus (or off-target viral protein) was spiked into a mixture of human saliva and extraction buffer to create the test sample, and then added to the sample pad before closing the folding card device to start the test. After 60 min, the test region was photographed. For comparison tests of commercial SARS-CoV-2 lateral flow assays, gamma-irradiated SARS-CoV-2 virus was added directly to the extraction buffer provided by the manufacturer.

### Viral RNA Detection

The nitrocellulose membrane and wicking pad were overlapped on an adherent backing material and cut into a 3-channel geometry with a laser cutter. The membrane was spotted with anti-DNA/RNA capture antibody in the test region and dried. Gamma-irradiated SARS-CoV-2 virus (or off-target synthetic viral RNA) was mixed with extraction buffer and the DNA probe set, heated on a heat block at 65 °C for 15 min, and loaded into a well (Channel 1) proximal to wells containing reagents for Channel 2 (DIG-labeled HCR hairpins h1 and h2) and Channel 3 (CB-labeled anti-DIG reporter antibody). The ends of the three membrane channels were simultaneously immersed into the three wells to start the test. After 90 min, the test region was photographed.

## ASSOCIATED CONTENT

### Supporting Information

Materials, additional methods, videos, replicate data, and additional studies.

## AUTHOR INFORMATION

### Authors

**Samuel J. Schulte** – *Division of Biology & Biological Engineering, California Institute of Technology, Pasadena, California 91125, United States;*

**Jining Huang** – *Division of Biology & Biological Engineering, California Institute of Technology, Pasadena, California 91125, United States;*

### Notes

The authors declare competing financial interests in the form of patents, pending patent applications, and the startup company Molecular Instruments.

## ACKNOWLEDGMENTS

We thank M. Schwarzkopf and K. S. Lee for performing preliminary studies. We thank M. E. Bronner for reading a draft of the manuscript. We thank M. Schwarzkopf, K. S. Lee, L. M. Hochrein, G. Shin, B. J. Wold, and R. F. Ismagilov for helpful discussions. We thank G. Shin of the Molecular Technologies resource within the Beckman Institute at Caltech for providing HCR reagents. This work was funded by the Shurl and Kay Curci Foundation, by the Richard N. Merkin Institute for Translational Research at Caltech, by the National Aeronautics and Space Administration (Translational Research Institute for Space Health; NNX16AO69A), and by the National Institutes of Health (NIBIB R01EB006192 and NIGMS training grant GM008042 to S.J.S.).

## ABBREVIATIONS USED

CB: carbon black
DIG: digoxigenin
HCR: hybridization chain reaction
LAMP: loop-mediated isothermal amplification
N: nucleocapsid
PR: polystreptavidin R
SARS-CoV-2: severe acute respiratory syndrome coronavirus 2.

