## Supplementary Information for "HCR Lateral Flow Assays for Amplified Instrument-Free At-Home SARS-CoV-2 Testing"

#### Contents

**S1 Materials and methods** **S3**
S1.1 Conjugation of CB-labeled anti-DIG reporter antibodies . . . S3
S1.2 Gel study of rapid HCR signal amplification . . . S3
S1.3 Viral protein detection . . . S4
S1.3.1 Modification of anti-N capture and signal antibodies . . . S4
S1.3.2 Lateral flow device assembly for viral protein detection . . . S4
S1.3.3 Performing a viral protein detection test . . . S5
S1.3.4 Testing commercial SARS-CoV-2 lateral flow assays . . . S6
S1.4 Viral RNA detection . . . S9
S1.4.1 DNA probe set purification . . . S9
S1.4.2 Lateral flow device assembly for viral RNA detection . . . S9
S1.4.3 Performing a viral RNA detection test . . . S10
S1.5 Measurement of HCR polymer length and amplification gain in the context of lateral flow assays . . . S11
S1.5.1 Measurement of HCR polymer length using Alexa647-labeled HCR hairpins . . . S11
S1.5.2 Measurement of HCR signal gain using DIG-labeled HCR hairpins and CB-labeled anti-DIG reporter antibodies . . . S11
S1.5.3 Quantitative image analysis . . . S11

**S2 Replicates for lateral flow assays** **S12**
S2.1 Replicates for viral protein detection: amplified HCR lateral flow assay . . . S12
S2.2 Replicates for commercial SARS-CoV-2 rapid antigen tests: unamplified lateral flow assays . . . S13
S2.3 Replicates for viral RNA detection: amplified HCR lateral flow assay . . . S15

**S3 Additional studies** **S16**
S3.1 Gel study of HCR polymerization at short time scales . . . S16
S3.2 Measurement of HCR polymer length and amplification gain in the context of lateral flow assays . . . S17
S3.3 Visualization of automated reagent delivery using a 3-channel lateral flow assay with food coloring . . . S19

#### List of Figures

S1 Nitrocellulose membrane and wicking pad dimensions for viral protein detection . . . S4
S2 Sample pad and conjugate pad dimensions for viral protein detection . . . S5
S3 Dimensions of the left page of the folding card device for viral protein detection . . . S6
S4 Dimensions of the right page of the folding card device for viral protein detection . . . S7
S5 Dimensions of the pressure bar for the folding card device used for viral protein detection . . . S7
S6 Steps for assembling the folding card device for viral protein detection . . . S8
S7 Nitrocellulose membrane and wicking pad dimensions for viral RNA detection . . . S9
S8 Photograph of the lateral flow device for viral RNA detection . . . S10
S9 Viral protein detection: sensitivity of amplified HCR lateral flow assay (cf. Figure 2a and Table 1) . . . S12
S10 Viral protein detection: background and cross-reactivity of amplified HCR lateral flow assay (cf. Figure 2bc) . . . S12
S11 Commercial SARS-CoV-2 rapid antigen test A: sensitivity of unamplified lateral flow assay (cf. Table 1) . . . S13

[#]Authors contributed equally. [†]Division of Biology & Biological Engineering, California Institute of Technology, Pasadena, CA 91125, USA. [‡]Division of Engineering & Applied Science, California Institute of Technology, Pasadena, CA 91125, USA. [*]

S12 Commercial SARS-CoV-2 rapid antigen test B: sensitivity of unamplified lateral flow assay (cf. Table 1) . . . . S13
S13 Commercial SARS-CoV-2 rapid antigen test C: sensitivity of unamplified lateral flow assay (cf. Table 1) . . . . S13
S14 Commercial SARS-CoV-2 rapid antigen test D: sensitivity of unamplified lateral flow assay (cf. Table 1) . . . . S13
S15 Commercial SARS-CoV-2 rapid antigen test E: sensitivity of unamplified lateral flow assay (cf. Table 1) . . . . S14
S16 Viral RNA detection: sensitivity of amplified HCR lateral flow assay (cf. Figure 4a) . . . . . . . . . . . . . . . S15
S17 Viral RNA detection: background and cross-reactivity of amplified HCR lateral flow assay (cf. Figure 4bc) . . S15
S18 Characterization of HCR polymerization at short time scales via agarose gel electrophoresis . . . . . . . . . . S16
S19 Measurement of HCR polymer length in the context of a lateral flow assay for viral protein detection . . . . . . S17
S20 Measurement of HCR amplification gain in the context of a lateral flow assay for viral protein detection . . . . S18
S21 Measurement of HCR amplification gain in the context of a lateral flow assay for viral RNA detection . . . . . S18

### List of Tables

S1 Estimated HCR polymer length in the context of a lateral flow assay for viral protein detection . . . . . . . . . S17
S2 Estimated amplification gain in the context of lateral flow assays for viral protein or RNA detection . . . . . . S18

### S1 Materials and methods

#### S1.1 Conjugation of CB-labeled anti-DIG reporter antibodies

An anti-digoxigenin (DIG) antibody (Jackson ImmunoResearch Laboratories, 200-002-156) was passively conjugated to Special Black 4 carbon black (CB; The Cary Company, 24W208).[1] A 30K Amicon Ultra-0.5 mL Centrifugal Filter (Millipore Sigma, UFC503096) was prepared by centrifuging 450 $\mu$L 5 mM borate buffer (Thermo Scientific, 28341, diluted 1:200 in nanopure water) at room temperature for 5 min at 13.8k rcf. To the prepared centrifugal filter, 200 $\mu$L anti-DIG antibody (1.3 mg/mL) was added and centrifuged at room temperature for 5 min at 13.8k rcf to buffer exchange the antibody. Then, 350 $\mu$L 5 mM borate buffer was added and centrifuged at room temperature for 5 min at 13.8k rcf. The 5 mM borate buffer wash was repeated four additional times. The column was then turned upside down in a new tube, and the buffer-exchanged antibody was recovered by centrifuging at room temperature for 5 min at 1k rcf. The concentration of the buffer-exchanged antibody was determined by absorbance at 280 nm on a NanoDrop spectrophotometer, and the concentration was normalized to 1 mg/mL by the addition of 5 mM borate buffer.

A 1% (w/v) carbon black solution was prepared fresh in nanopure water, and the entire volume (2 mL) was continuously sonicated (cycle 1.0) for 15 min at room temperature at 100% amplitude with an ultrasonic processor (Hielscher Ultrasound Technology, UP50H). To 200 $\mu$L of 1 mg/mL buffer-exchanged anti-DIG antibody, 570 $\mu$L 0.2% carbon black (diluted 1:5 in 5 mM borate buffer) was added in a SuperSpin Polypropylene tube (Thomas Scientific, 20A00L069). The solution was pipetted up and down before mixing end-over-end on a rotary wheel for 3 h at room temperature. A 1% (w/v) BSA (Sigma-Aldrich, A2153) in 5 mM borate buffer solution was prepared, and the pH was adjusted to 8.5 by adding 1 M sodium hydroxide (NaOH) (Sigma-Aldrich, S2770). After 3 h of end-over-end rotation, 700 $\mu$L 1% (w/v) BSA in 5 mM borate buffer (pH 8.5) was added to the carbon black mixture, and the mixture was vortexed briefly before end-over-end mixing on a rotary wheel for 15 min at room temperature to quench any unmodified carbon black. The mixture was centrifuged at room temperature for 15 min at 13.8k rcf, and the supernatant was discarded. A total of four times: the carbon black antibody conjugate was resuspended in 1 mL 1% (w/v) BSA in 5 mM borate buffer (pH 8.5) by pipetting up and down and vortexing, the mixture was centrifuged at room temperature for 15 min at 13.8k rcf, and the supernatant was discarded. The CB-labeled anti-DIG antibodies were lastly resuspended in 1% (w/v) BSA in 5 mM borate buffer (pH 8.5) with 0.02% sodium azide (Sigma-Aldrich, S2002). The CB-labeled anti-DIG antibodies were stored at 4 °C.

#### S1.2 Gel study of rapid HCR signal amplification

DNA HCR hairpins h1 and h2 at 3 $\mu$M (Molecular Technologies, B3-Alexa647) were separately snap-cooled (with h1 and h2 in separate tubes) by heating to 95 °C for 90 sec and cooling at room temperature in a dark drawer for at least 30 min. HPLC-purified HCR initiator i1 (Molecular Technologies, B3) was diluted in IDTE (pH 8.0) (Integrated DNA Technologies, 11-05-01-09) to 0.03 $\mu$M (0.01× initiator:hairpin ratio reaction). To a new tube was added: 1.2 $\mu$L 5× saline sodium citrate (SSC) (Life Technologies, 15557-044) with 1% Tween-20 (Teknova, T0025), 4.8 $\mu$L 5× SSC, 2 $\mu$L snap-cooled hairpin h1, and 2 $\mu$L snap-cooled hairpin h2. To trigger hairpin polymerization, 2 $\mu$L of diluted i1 was added and allowed to react for 10 min. For the leakage lane, initiator i1 was omitted, and an additional 2 $\mu$L IDTE (pH 8.0) was added and allowed to incubate for 10 min. To analyze HCR polymer formation, 2.4 $\mu$L 6× DNA Gel Loading Dye (Thermo Scientific, R0611) was added, and 12 $\mu$L of the reaction mixture was loaded into a 4.8 mm-wide well of a 1% (w/v) agarose (Invitrogen, 16500500) gel cast and run in 1× lithium borate (LB) buffer (Faster Better Media, LB20-10). A dsDNA ladder pre-stained with SYBR Gold was also loaded into the gel. To stain the ladder, a loading dye solution with SYBR Gold (Invitrogen, S11494) was first created by adding 1 $\mu$L 10,000× SYBR Gold solution to 400 $\mu$L 6× DNA Gel Loading Dye (Thermo Scientific, R0611). Then, to 11.04 $\mu$L nanopure water, 0.96 $\mu$L of GeneRuler 1 kb Plus DNA Ladder (Thermo Scientific, SM1331) and 2.40 $\mu$L loading dye solution with SYBR Gold were added, and 12 $\mu$L of this solution was loaded into the gel. The gel was run for 40 min at 150 V before imaging with an Amersham ImageQuant 800 Fluor imaging system (GE Life Sciences, 29399484) with the Cy2 (to image SYBR Gold) and Cy5 (to image Alexa647) filters.

#### S1.3 Viral protein detection

##### S1.3.1 Modification of anti-N capture and signal antibodies

The anti-N capture antibody (Sino Biological, 40143-MM08) was conjugated to NHS-dPEG$_{12}$-biotin (Quanta Biodesign Limited, 10198).[2] A 30k Amicon Ultra-0.5 mL Centrifugal Filter (Millipore Sigma, UFC503096) was prepared by centrifuging 500 $\mu$L 1× phosphate-buffered saline (PBS) (Ambion, AM9624) at room temperature for 5 min at 13.8k rcf. In a separate tube, 50 $\mu$L anti-N antibody (1 mg/mL) was combined with 400 $\mu$L 1× PBS, and the entire solution was loaded into the centrifugal column and centrifuged at room temperature for 5 min at 13.8k rcf to buffer exchange the antibody. Two times, 350 $\mu$L 1× PBS was added and centrifuged at room temperature for 5 min at 13.8k rcf. The column was then turned upside down in a new tube, and the buffer-exchanged antibody was recovered by centrifuging at room temperature for 5 min at 1k rcf. The concentration of the buffer-exchanged antibody was determined by absorbance at 280 nm on a NanoDrop spectrophotometer. To 10 mg NHS-dPEG$_{12}$-biotin, 800 $\mu$L 1× PBS was added, and this solution was added to the buffer-exchanged antibody at a 50:1 molar ratio (NHS-dPEG$_{12}$-biotin:buffer-exchanged antibody). After incubating the antibody biotinylation reaction at room temperature for 1 h, 500 $\mu$L 1× PBS was centrifuged at room temperature for 5 min at 13.8k rcf in a new centrifugal filter. To the biotinylated antibody solution, 400 $\mu$L 1× PBS was added, and the entire volume was loaded into the centrifugal filter and centrifuged at room temperature for 5 min at 13.8k rcf. A total of four times: 350 $\mu$L 1× PBS was added and centrifuged at room temperature for 5 min at 13.8k rcf. The column was turned upside down in a new tube, and the biotinylated antibody was recovered by centrifuging at room temperature for 5 min at 1k rcf. The concentration of the biotinylated antibody was determined by absorbance at 280 nm on a NanoDrop spectrophotometer, and the biotinylated antibody was diluted to 1 mg/mL with 1× PBS, aliquoted, and stored at -20 °C.

The anti-N signal antibody (Sino Biological, 40143-MM05) was conjugated to HCR initiator i1 (Molecular Technologies, B3) with an Antibody-Oligonucleotide All-in-One Conjugation Kit (Vector Laboratories, A-9202-001) according to the manufacturer's instructions. The concentration of the initiator-labeled antibody was determined with a Pierce BCA Protein Assay Kit (Thermo Scientific, 23225).

##### S1.3.2 Lateral flow device assembly for viral protein detection

A 25 mm backed nitrocellulose membrane (Cytiva, FF80HP PLUS grade, 10547042) was adhered onto ultra optically clear double-sided tape (McMaster-Carr, 90727A110). A 30 mm wicking pad (Cytiva, CF7 grade, 8117-6621) was also adhered onto the double-sided tape with a 5 mm overlap on top of the nitrocellulose membrane. The wicking pad and nitrocellulose were then cut to size with a laser cutter (Full Spectrum Laser, Muse Core Desktop $CO_2$ Laser Cutter; see Figure S1 for dimensions).

The sample pad (Ahlstrom-Munksjö, Chopped Glass w/Binder, Grade 8951) and conjugate pads (Ahlstrom-Munksjö, Chopped Glass w/Binder, Grade 8964) were cut to size with a laser cutter (see Figure S2 for dimensions), submerged in blocking solution for 15 min with end-over-end rotation on a rotary wheel, and dried flat for 1 h in a fan-equipped oven (Sci-Gene Model 2000 Micro Hybridization Incubator, 1040-50-1) set to 37 °C. The sample pad blocking solution consisted of 0.2% Blocker bovine serum albumin (BSA) (Thermo Fisher Scientific, 37525) and 0.2% Tween-20 (Teknova, T0025). The

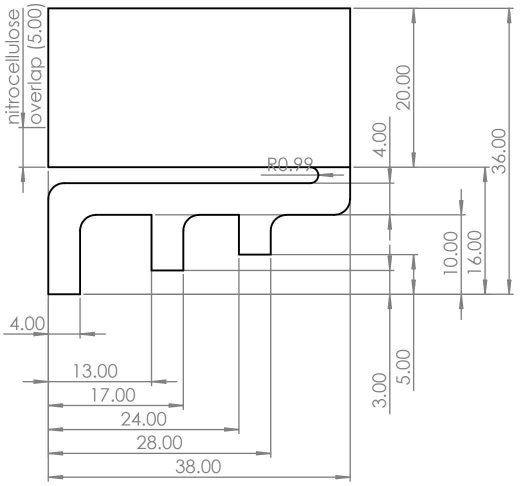


**Figure S1: Nitrocellulose membrane and wicking pad dimensions for viral protein detection.** The nitrocellulose membrane and wicking pad were overlapped 5 mm on ultra optically clear double-sided tape and cut to size with a laser cutter. All dimensions are shown in millimeters (R: radius). All curved lines have a radius of 2.00 mm unless otherwise indicated.

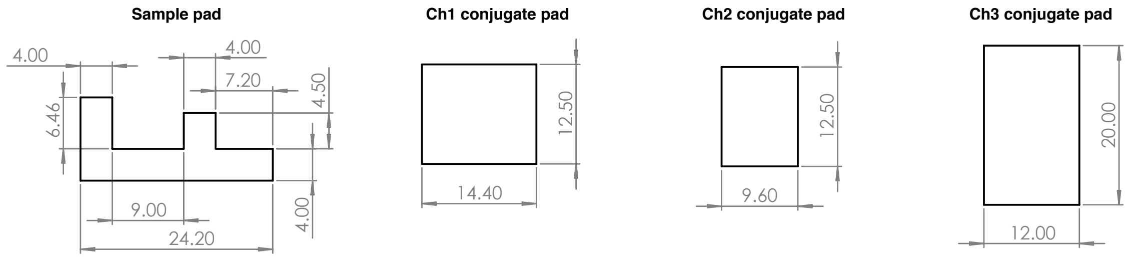


**Figure S2: Sample pad and conjugate pad dimensions for viral protein detection.** Pads were cut to size with a laser cutter. All dimensions are shown in millimeters.

Channel 1 conjugate pad blocking solution consisted of 1% Blocker BSA, 1% sucrose (Sigma-Aldrich, S7903), 1% trehalose (Sigma-Aldrich, T9531), 0.02% Tween-20, and 0.2× PBS. The Channel 2 conjugate pad blocking solution consisted of 0.5% BSA (Sigma-Aldrich, A2153), 0.2% sucrose, 0.2% trehalose, 0.1% Tween-20, 1× SSC, and 0.1% dextran sulfate sodium salt (Sigma-Aldrich, D6001). The Channel 3 conjugate pad blocking solution consisted of 0.5% BSA, 0.2% sucrose, 0.5% Tween-20, and 0.2× PBS.

For each Channel 1 conjugate pad: to 75 $\mu$L Channel 1 conjugate pad blocking solution, biotinylated anti-N capture antibody was added to 2 $\mu$g/mL, and initiator-labeled anti-N signal antibody was added to 0.125 $\mu$g/mL. This solution was mixed by pipetting up and down, and 75 $\mu$L of this solution was pipetted onto the blocked Channel 1 conjugate pad. For each Channel 2 conjugate pad: 4 $\mu$L each of DIG-labeled HCR hairpins h1 and h2 (Molecular Instruments; B3-DIG) were separately snap-cooled (with h1 and h2 in separate tubes) by heating to 95 °C for 90 sec followed by cooling at room temperature for 30 min. To 50 $\mu$L Channel 2 conjugate pad blocking solution, 4 $\mu$L of each snap-cooled DIG-labeled HCR hairpin (h1 and h2) was added. For h1-only assays run without HCR hairpin h2, 4 $\mu$L HCR hairpin buffer was instead added (Molecular Technologies). This solution was mixed by pipetting up and down, and 50 $\mu$L of this solution was pipetted onto the blocked Channel 2 conjugate pad. For each Channel 3 conjugate pad: 4 $\mu$L CB-labeled anti-DIG reporter antibody was added to a microcentrifuge tube on ice and continuously sonicated (cycle 1.0) for 60 sec at 40% amplitude with an ultrasonic processor (Hielscher Ultrasound Technology, UP50H). To 115 $\mu$L Channel 3 conjugate pad blocking solution, 4 $\mu$L freshly sonicated CB-labeled anti-DIG reporter antibody was added. This solution was mixed by pipetting up and down, and 115 $\mu$L of this solution was pipetted onto the blocked Channel 3 conjugate pad. All conjugate pads were then dried flat for 1 h in a fan-equipped oven (SciGene Model 2000 Micro Hybridization Incubator, 1040-50-1) set to 25 °C.

A folding card device was created from 1.75 mm white polylactic acid (PLA) filament (HATCHBOX) with a 3D printer (Creality, Ender-5 Plus 3D Printer) with a 200 °C nozzle temperature and 60 °C bed temperature (see Figures S3–S5 for dimensions of the left page, right page, and pressure bar). Magnets (McMaster-Carr, 5862K102) were affixed into the device with hot glue to aid in holding the left and right pages together once the card is closed. Double-sided tape (McMaster-Carr, 7602A58) was cut with a laser cutter to the dimensions of the sample pad pedestal and applied to the sample pad pedestal. Double-sided tape was cut to shape with a laser cutter and applied to the conjugate pad pedestals, in each case leaving an exposed 2 mm region of the conjugate pad pedestal where the sample pad extends onto the conjugate pad pedestals. The adhesive backing was removed from the double-sided tape on the sample pad pedestal and conjugate pad pedestals, and the blocked sample pad was adhered to the sample pad pedestal, with the ends of each sample pad channel extending onto the conjugate pad pedestals. The conjugate pads were adhered to the double-sided tape on the conjugate pad pedestals, with the leading edge of the conjugate pads overlapping on top of the three sample pad channels.

Polystreptavidin R (Biotez, 10120030) was diluted to 0.94 mg/mL with 1× PBS. At the test region on the nitrocellulose membrane, 0.55 $\mu$L diluted polystreptavidin R was gently spotted with a P2 pipette. The nitrocellulose membrane was allowed to dry at room temperature for 30 min. Then, the adhesive backing was removed from the nitrocellulose and wicking pad, and the nitrocellulose and wicking pad were adhered to the left page of the folding card device. The pressure bar was then put in place on the left page of the device to apply pressure to the junction of the wicking pad and nitrocellulose membrane. The left and right pages of the folding card device were then assembled at the hinge (see Figure S6).

##### S1.3.3 Performing a viral protein detection test

Pooled human saliva collected before November 2019 (Lee Biosolutions, 991-05-P-PreC) was thawed at room temperature. Gamma-irradiated SARS-CoV-2 (BEI Resources, NR-52287), recombinant OC43 N protein (Sino Biological, 40643-V07E), or recombinant Influenza A H3N2 nucleoprotein (Sino Biological, 40499-V08B), was diluted in 1× PBS as needed, and 1 $\mu$L of the diluted solution was added to 299 $\mu$L of a solution of 2/3 saliva and 1/3 extraction buffer (5× SSC with 0.1% Tween-20). Then, 300 $\mu$L was slowly applied to the circular region of the sample pad pedestal with a pipette, and the top of the folding card device was closed to start the test. After 60 min, the device was placed in a 17-inch light tent (Angler, CT-DSLEDII) at maximum light intensity with a black background and photographed with a camera (Panasonic GH4) equipped with a 60 mm macro lens (Olympus, V312010BU000) with the following settings: 1/2000 sec shutter speed, 200 ISO, f/2.8 aperture, and neutral white balance.

##### S1.3.4 Testing commercial SARS-CoV-2 lateral flow assays

Gamma-irradiated SARS-CoV-2 (BEI Resources, NR-52287) was diluted in 1× PBS as needed, and 1 $\mu$L of the diluted virus solution was added directly to the commercial test extraction buffer to reach the desired final concentration. This solution was then added to the device according to the manufacturers' instructions. After the minimum manufacture-recommended test time had elapsed, the device was photographed as described above.

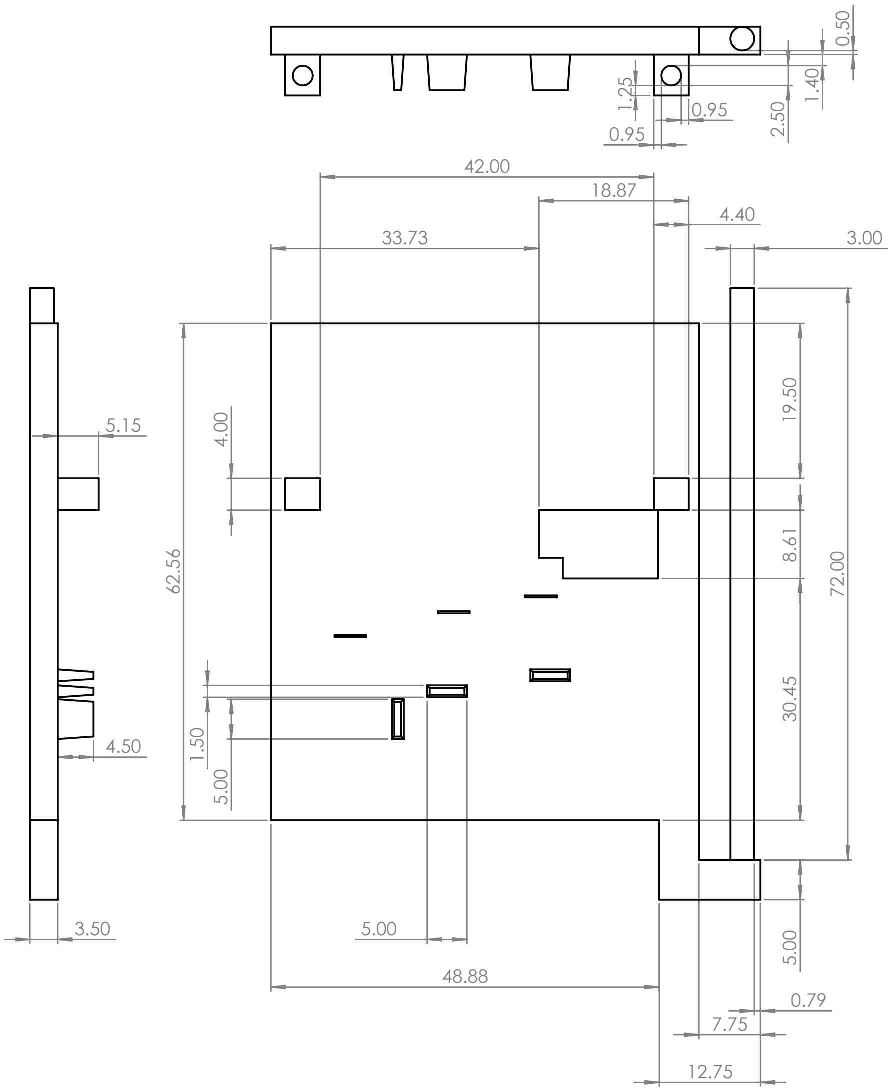


**Figure S3: Dimensions of the left page of the folding card device for viral protein detection.** The left page of the folding card device was 3D-printed. All dimensions are shown in millimeters.

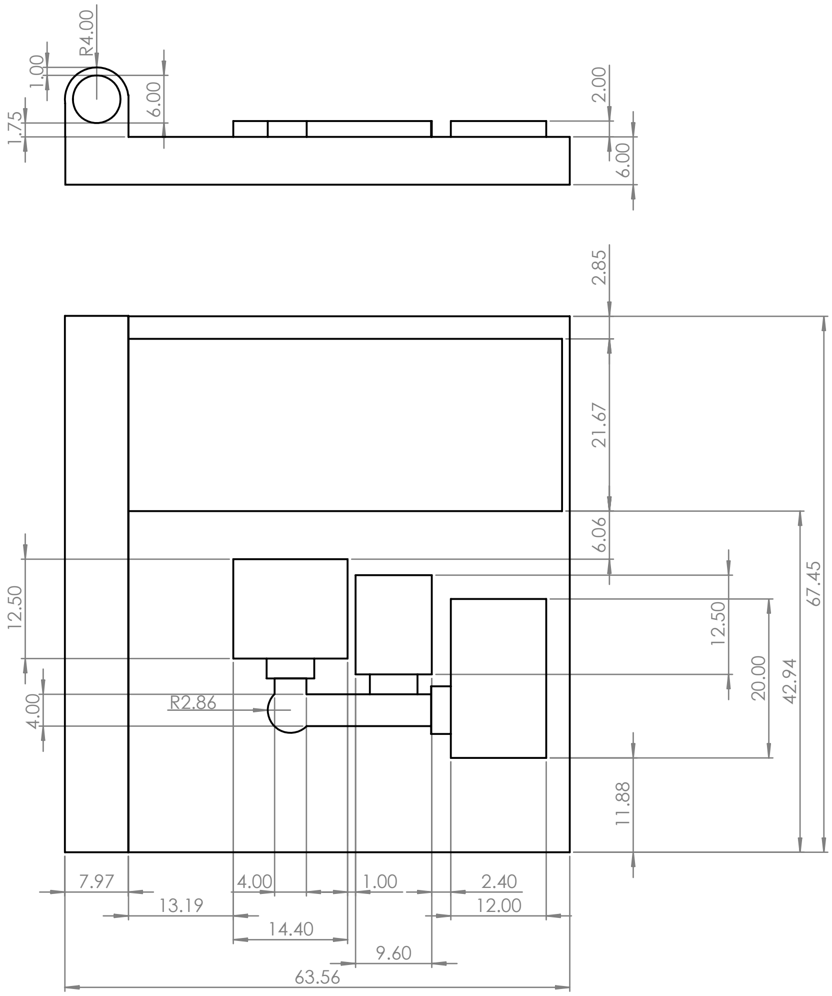


**Figure S4: Dimensions of the right page of the folding card device for viral protein detection.** The right page of the folding card device was 3D-printed. All dimensions are shown in millimeters (R: radius).

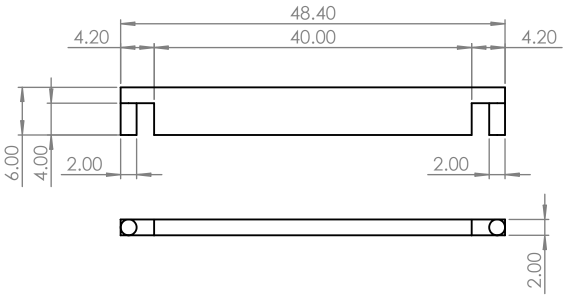


**Figure S5: Dimensions of the pressure bar for the folding card device used for viral protein detection.** The pressure bar was 3D-printed and attached to the left page of the folding card device to create pressure between the wicking pad and nitrocellulose membrane. All dimensions are shown in millimeters.

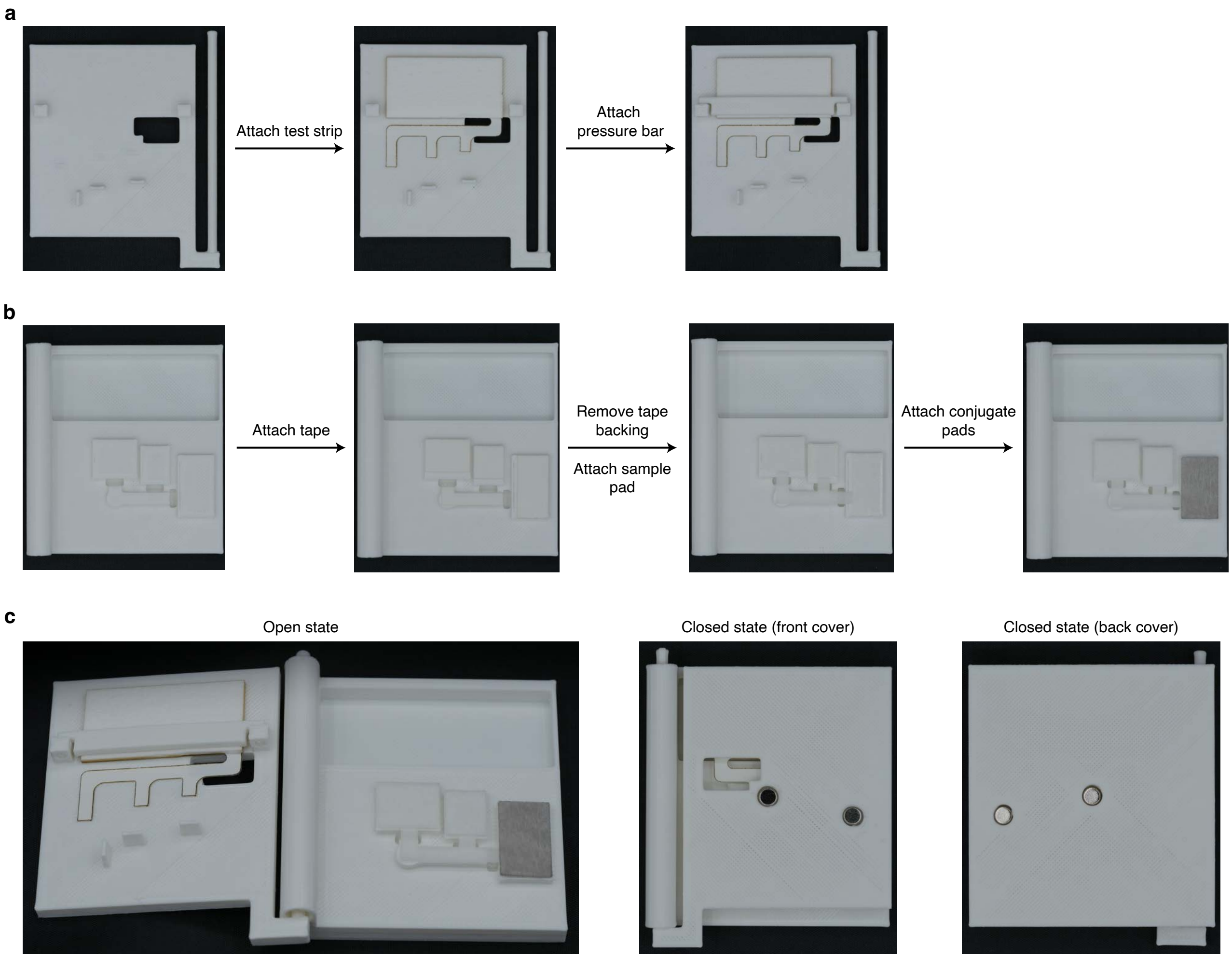


**Figure S6: Steps for assembling the folding card device for viral protein detection.** (a) Assembling the left page. The adherent backing on the nitrocellulose membrane and wicking pad is removed, and the bottoms of the three nitrocellulose membrane channels are aligned with the thin markers on the device before adhering the nitrocellulose membrane and wicking pad in place. The pressure bar is added by pushing its two cylindrical prongs into the holes on either side of the wicking pad. (b) Assembling the right page. Double-sided tape is applied to the sample pad and conjugate pad pedestals. The adherent backing is removed from the double sided tape, and the sample pad is affixed to the sample pad pedestal. The conjugate pads are then applied to the conjugate pad pedestals, with their leading edges overlapping with the sample pad by 2 mm. (c) The left and right pages of the folding card device are assembled at the hinge. Images depict the device in the open state (left) and in the closed state viewing either the front cover (middle) or back cover (right).

#### S1.4 Viral RNA detection

##### S1.4.1 DNA probe set purification

All probes in the SARS-CoV-2 DNA probe set (Molecular Technologies, 4014/E1052) were pooled and purified with polyacrylamide gel electrophoresis (PAGE). A 1.5 mm 20% denaturing polyacrylamide 19 cm $\times$ 20 cm gel was prepared and run in 1$\times$ tris-borate-ethylenediaminetetraacetic acid (EDTA) (TBE) (VWR, 75800-954). The gel was pre-run at 400 V for 30 min to warm the gel. The DNA probes (1000 $\mu$L, with each probe at 0.5 $\mu$M) were mixed 1:1 with denaturing loading buffer consisting of 88.8% formamide (Ambion, AM9342), 10 mM EDTA (Invitrogen, 15575-038), 2 mM Tris-HCl (pH 7.5) (Invitrogen, 15567027), 62.5 $\mu$g/mL bromophenol blue (MP Biomedicals, 193990), and 62.5 $\mu$g/mL xylene cyanol (Bio-Rad, 161-0423), and heated to 95 °C for 5 min. The entire mixture was then immediately loaded into a single well and run at 400 V for 3.5 h. The gel was unmolded onto a plastic-wrapped chromatography plate, and the DNA probe band was cut from the gel under a short wavelength UV shadow, sliced into 0.5 cm $\times$ 0.5 cm pieces, and placed in a 50 mL LoBind tube (Eppendorf, 0030122232). To the gel slices, 15 mL of 0.3 M NaCl (Promega, V4221) was added, and the tube was inverted end-over-end on a rotary wheel overnight ($>$ 12 h). The next day, a sterile flip filter (Millipore, SCGP00525) was used to filter the liquid into a new 50 mL LoBind tube. To the filtered liquid, 35 mL 100% ethanol (Koptec, V1001) and 20 $\mu$L molecular biology grade glycogen (Thermo Scientific, R0561) were added, and the mixture was inverted several times to mix before incubating overnight at $-20$ °C. The 50 mL tube was then centrifuged at 4 °C for 30 min at 4.5k rcf. The supernatant was carefully removed with a pipet and discarded, and any remaining liquid was allowed to evaporate for 5 h with a Kimwipe covering the top of the tube. The pellet was resuspended in 500 $\mu$L IDTE (pH 8.0), and the total probe concentration was determined by 260 nm absorbance with a NanoDrop spectrophotometer.

##### S1.4.2 Lateral flow device assembly for viral RNA detection

A 25 mm backed nitrocellulose membrane (Cytiva, FF80HP PLUS grade, 10547042) was adhered onto a polyethylene backing card (McMaster-Carr, 1441T52). A 25 mm wicking pad (Cytiva, CF7 grade, 8117-6621) was also adhered onto the backing card with a 2 mm overlap on top of the nitrocellulose membrane. The wicking pad and nitrocellulose were then cut to size with a laser cutter (see Figure S7 for dimensions and Figure S8 for a photograph of the device). With a P2 pipette, 0.70 $\mu$L of an anti-DNA:RNA capture antibody (0.8 mg/mL) (Abcam, ab256361) was gently spotted at the test region of the nitrocellulose membrane. The nitrocellulose membrane was allowed to dry at room temperature for at least 30 min. A paper clip was added to apply pressure to the junction of the wicking pad and the nitrocellulose membrane (see Figure S8).

For each assay, to prepare the Channel 1 solution, 540 pmol of split-initiator DNA probes were added to 100 $\mu$L of Channel 1 buffer (created by combining the following reagents in nanopure water in the listed order to reach the indicated concentration: 1.5% BSA, 4$\times$ SSC, 0.05% Tween-20, 0.4 U/$\mu$L RNase inhibitor (Promega, N2615), and 0.2% dextran sulfate sodium salt). For each assay, to prepare the Channel 2 solution, 4 $\mu$L each of DIG-labeled HCR hairpins h1 and h2 (Molecular Instruments; B3-DIG) were separately snap-cooled (with h1 and h2 in separate tubes) by heating to 95 °C for 90 sec followed by cooling at room temperature for 30 min. To 92 $\mu$L Channel 2 buffer, which consisted of 0.2% dextran sulfate sodium salt in 5$\times$ SSC

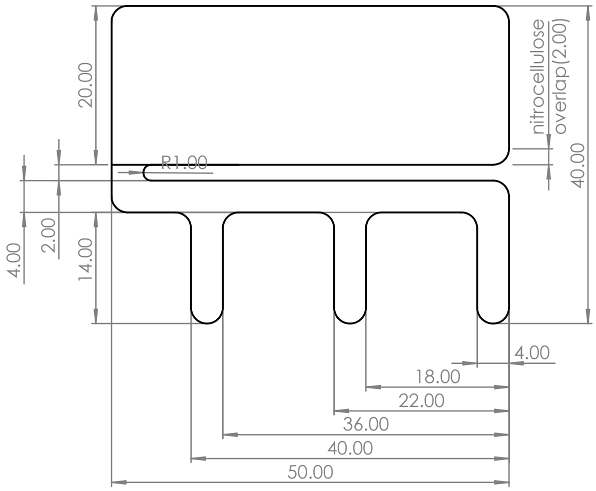


**Figure S7: Nitrocellulose membrane and wicking pad dimensions for viral RNA detection.** The nitrocellulose membrane and wicking pad were overlapped on a polyethylene backing card and cut to size with a laser cutter. All dimensions are shown in millimeters (R: radius). All curved lines have a radius of 2.00 mm unless otherwise indicated.

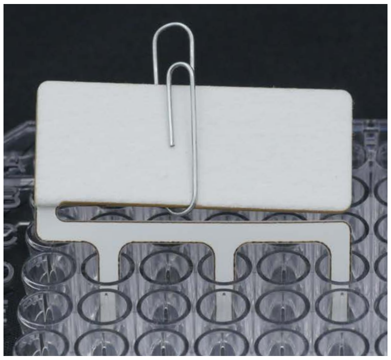

**Figure S8: Photograph of the lateral flow device for viral RNA detection.** The lateral flow device with three channels resting within their respective wells on a 96-well plate.

with 0.1% Tween-20, 4 $\mu$L each of snap-cooled DIG-labeled HCR hairpins h1 and h2 were added, and the solution was mixed by pipetting up and down and placed in a well (Channel 2 well) of a 96-well plate (Greiner Bio-One, 655901). For h1-only assays run without HCR hairpin h2, 4 $\mu$L HCR hairpin buffer was instead added (Molecular Technologies). For each assay, to prepare the Channel 3 solution, 4 $\mu$L CB-labeled anti-DIG reporter antibody was added to a microcentrifuge tube on ice and continuously sonicated (cycle 1.0) for 60 sec at 40% amplitude with an ultrasonic processor. To 96 $\mu$L Channel 3 buffer (created by combining the following reagents in nanopure water in the listed order to reach the indicated concentration: 0.35% BSA, 5$\times$ SSC, and 0.1% Tween-20), 4 $\mu$L freshly sonicated CB-labeled anti-DIG reporter antibody was added, and the solution was mixed by pipetting up and down and placed in a well of a 96-well plate (Channel 3 well) right-adjacent to the Channel 2 well.

##### S1.4.3 Performing a viral RNA detection test

Gamma-irradiated SARS-CoV-2 (BEI Resources, NR-52287) or synthetic genomic RNA from 229E (American Type Culture Collection, VR-740D) or HKU1 (American Type Culture Collection, VR-3262SD) coronaviruses was diluted as needed in nanopure water and added to the Channel 1 solution and mixed by pipetting up and down. This solution was incubated at 65 °C for 15 min on a heat block and transferred to a well of a 96-well plate (Channel 1 well) left-adjacent to the Channel 2 well. To start the lateral flow assay, the ends of the three nitrocellulose channels (Channel 1, Channel 2, Channel 3) were simultaneously submerged in their respective wells in the 96-well plate. After 90 min, the test strip was placed in a 17-inch light tent (Angler, CT-DSLEDII) at maximum light intensity with a white background and photographed with a camera (Panasonic GH4) equipped with a 60 mm macro lens (Olympus, V312010BU000) with the following settings: 1/4000 sec shutter speed, 200 ISO, f/2.8 aperture, and neutral white balance.

#### S1.5 Measurement of HCR polymer length and amplification gain in the context of lateral flow assays

##### S1.5.1 Measurement of HCR polymer length using Alexa647-labeled HCR hairpins

To measure the average HCR polymer length, we do $N = 3$ replicate lateral flow assays for each of two types of experiment:

- **Amplified experiment (h1 and h2):** using both HCR hairpins (Alexa647-labeled h1 and Alexa647-labeled h2) in Channel 2 so that each HCR initiator labeling an anti-N antibody captured at the test region can trigger polymerization of a tethered Alexa647-decorated amplification polymer.
- **Unamplified experiment (h1 only):** use only HCR hairpin h1 (Alexa647-labeled h1) so that HCR polymerization cannot proceed and each HCR initiator labeling an anti-N antibody captured at the test region can bind only one Alexa647-labeled h1 hairpin.

To avoid interference with the fluorescent hairpin signal, the CB-labeled anti-DIG reporter antibody was omitted from the Channel 3 conjugate pad solution. Gamma-irradiated SARS-CoV-2 was spiked into a mixture of saliva and extraction buffer to a final concentration of 20,000 copies/$\mu$L for all tests. After 1 h, the test strip was removed from the folding card device, and fluorescence from the Alexa647-labeled hairpins was imaged with an FLA-5100 fluorescent scanner (Fujifilm Life Science) via a 635 nm laser and 665 nm long-pass filter. The mean HCR polymer length was then calculated as the ratio of amplified to unamplified intensities as described in Section S1.5.3.

##### S1.5.2 Measurement of HCR signal gain using DIG-labeled HCR hairpins and CB-labeled anti-DIG reporter antibodies

To calculate the HCR amplification gain, we do $N = 3$ replicate lateral flow assays for each of two types of experiment:

- **Amplified experiment (h1 and h2):** using both HCR hairpins (DIG-labeled h1 and DIG-labeled h2) in Channel 2 so that each HCR initiator labeling an anti-N antibody captured at the test region can trigger polymerization of a tethered DIG-decorated amplification polymer, which can then be bound by multiple CB-labeled anti-DIG reporter antibodies from Channel 3.
- **Unamplified experiment (h1 only):** use only HCR hairpin h1 (DIG-labeled h1) so that HCR polymerization cannot proceed and each HCR initiator labeling an anti-N antibody captured at the test region can bind only one DIG-labeled h1 hairpin, which can then be bound by one CB-labeled anti-DIG reporter antibody from Channel 3.

For viral protein detection tests, gamma-irradiated SARS-CoV-2 was spiked into a mixture of saliva and extraction buffer to a final concentration of 5,000 copies/$\mu$L and the test was run and photographed according to Section S1.3.3. For viral RNA detection tests, gamma-irradiated SARS-CoV-2 was spiked into extraction buffer to a final concentration of 5,000 copies/$\mu$L and the test was run and photographed according to Section S1.4.3. Image were converted to grayscale and the HCR amplification gain was then calculated as the ratio of amplified to unamplified intensities as described in Section S1.5.3. Ideally, the amplification gain would match the mean polymer length, corresponding to a situation in which a reporter antibody is binding to each hairpin within an amplification polymer.

##### S1.5.3 Quantitative image analysis

For each replicate of an amplified or unamplified experiment, the background-subtracted signal is calculated by taking the mean intensity in a signal box surrounding the test region and subtracting the mean intensity over one or two adjacent background boxes containing the same total number of pixels as the signal box. Let $\bar{x}_{\mathrm{h1+h2}}$ and $s_{\mathrm{h1+h2}}$ denote the sample mean and standard error of the mean for the background-subtracted amplified replicates, and let $\bar{x}_{\mathrm{h1}}$ and $s_{\mathrm{h1}}$ denote the sample mean and standard error of the mean for the background-subtracted unamplified replicates. The ratio of amplified to unamplified performance is then calculated as:

$$\bar{x}_{\mathrm{ratio}} = \bar{x}_{\mathrm{h1+h2}} / \bar{x}_{\mathrm{h1}} \tag{S1}$$

with standard error estimated via uncertainty propagation as

$$s_{\mathrm{ratio}} \leq \frac{\bar{x}_{\mathrm{h1+h2}}}{\bar{x}_{\mathrm{h1}}} \sqrt{\left(\frac{s_{\mathrm{h1+h2}}}{\bar{x}_{\mathrm{h1+h2}}}\right)^2 + \left(\frac{s_{\mathrm{h1}}}{\bar{x}_{\mathrm{h1}}}\right)^2}. \tag{S2}$$

This upper bound on estimated standard error holds under the assumption that the correlation between amplified and unamplified intensity is non-negative.

### S2 Replicates for lateral flow assays

#### S2.1 Replicates for viral protein detection: amplified HCR lateral flow assay

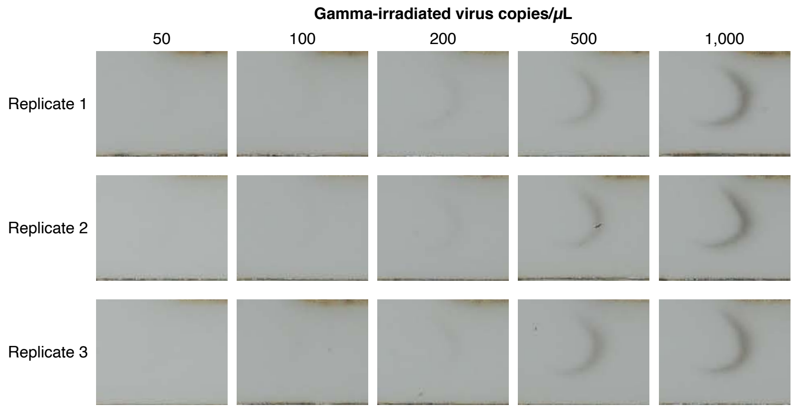


**Figure S9: Viral protein detection: sensitivity of amplified HCR lateral flow assay (cf. Figure 2a and Table 1).** Following the methods of Section S1.3.3, gamma-irradiated SARS-CoV-2 was spiked into a mixture of saliva and extraction buffer at the target concentration and loaded onto the sample pad before closing the folding card device to start the test. The test region was photographed after 60 minutes. $N = 3$ replicate assays at each target concentration. The test region is visible in all three replicates down to a limit of detection of 200 copies/$\mu$L.

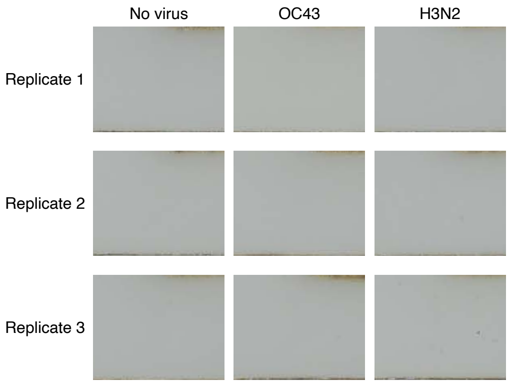


**Figure S10: Viral protein detection: background and cross-reactivity of amplified HCR lateral flow assay (cf. Figure 2bc).** Following the methods of Section S1.3.3, no virus (to measure background) or off-target recombinant viral OC43 N protein (83.74 ng/mL) or Influenza A (H3N2) (50.43 ng/mL) nucleoprotein (to test cross-reactivity) were spiked into a mixture of saliva and extraction buffer and loaded onto the sample pad before closing the folding card device to start the test. For the cross-reactivity tests, the off-target viral proteins were spiked in at high concentrations equivalent to $\approx 10^6$ virions/$\mu$L.[3,4] The test region was photographed after 60 min. $N = 3$ replicate assays at each target concentration. For each target type, no staining was visible at the test region for all three replicates, indicating that there is no visible background when SARS-CoV-2 is absent, and no visible cross-reactivity with off-target OC43 or H3N2 proteins.

#### S2.2 Replicates for commercial SARS-CoV-2 rapid antigen tests: unamplified lateral flow assays

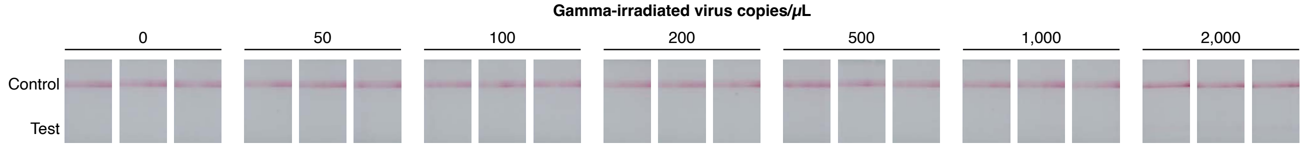


**Figure S11: Commercial SARS-CoV-2 rapid antigen test A: sensitivity of unamplified lateral flow assay (cf. Table 1).** Following the methods of Section S1.3.4, gamma-irradiated SARS-CoV-2 was spiked into the extraction buffer provided by the test manufacturer and loaded onto the commercial test device per the manufacturer's instructions. The test region was photographed after 15 min. $N = 3$ replicate assays at each target concentration. The test region is visible in all three replicates down to a limit of detection of 2000 copies/$\mu$L.

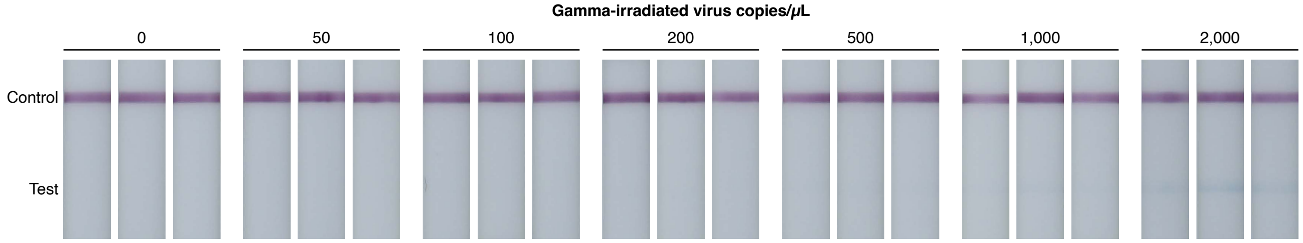


**Figure S12: Commercial SARS-CoV-2 rapid antigen test B: sensitivity of unamplified lateral flow assay (cf. Table 1).** Following the methods of Section S1.3.4, gamma-irradiated SARS-CoV-2 was spiked into the extraction buffer provided by the test manufacturer and loaded onto the commercial test device per the manufacturer's instructions. The test region was photographed after 10 min. $N = 3$ replicate assays at each target concentration. The test region is visible in all three replicates down to a limit of detection of 2000 copies/$\mu$L.

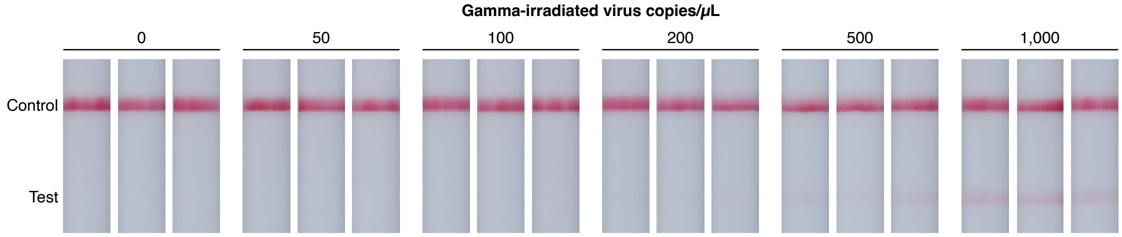


**Figure S13: Commercial SARS-CoV-2 rapid antigen test C: sensitivity of unamplified lateral flow assay (cf. Table 1).** Following the methods of Section S1.3.4, gamma-irradiated SARS-CoV-2 was spiked into the extraction buffer provided by the test manufacturer and loaded onto the commercial test device per the manufacturer's instructions. The test region was photographed after 15 min. $N = 3$ replicate assays at each target concentration. The test region is visible in all three replicates down to a limit of detection of 500 copies/$\mu$L.

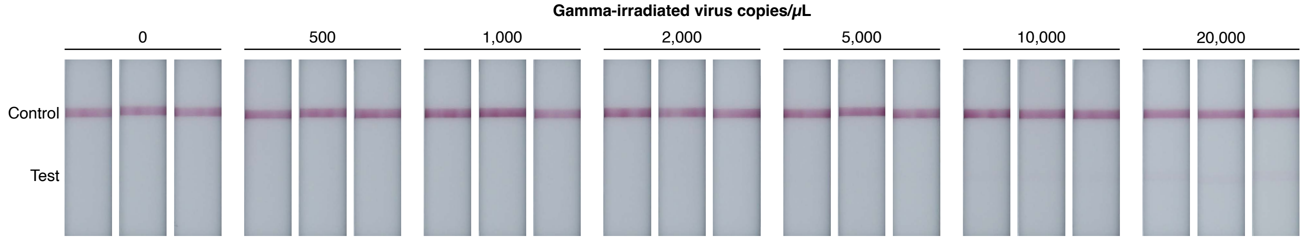


**Figure S14: Commercial SARS-CoV-2 rapid antigen test D: sensitivity of unamplified lateral flow assay (cf. Table 1).** Following the methods of Section S1.3.4, gamma-irradiated SARS-CoV-2 was spiked into the extraction buffer provided by the test manufacturer and loaded onto the commercial test device per the manufacturer's instructions. The test region was photographed after 15 min. $N = 3$ replicate assays at each target concentration. The test region is visible in all three replicates down to a limit of detection of 20,000 copies/$\mu$L.

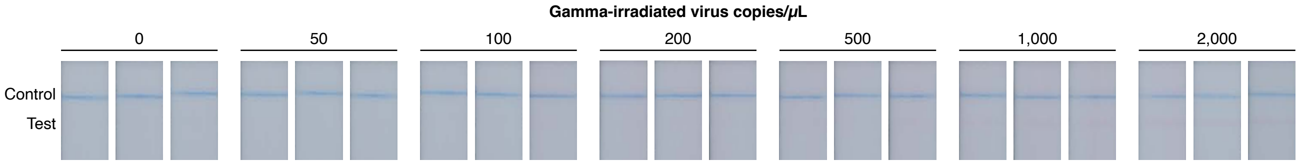


**Figure S15: Commercial SARS-CoV-2 rapid antigen test E: sensitivity of unamplified lateral flow assay (cf. Table 1).** Following the methods of Section S1.3.4, gamma-irradiated SARS-CoV-2 was spiked into the extraction buffer provided by the test manufacturer and loaded onto the commercial test device per the manufacturer's instructions. The test region was photographed after 10 min. $N = 3$ replicate assays at each target concentration. The test region is visible in all three replicates down to a limit of detection of 1000 copies/$\mu$L.

#### S2.3 Replicates for viral RNA detection: amplified HCR lateral flow assay

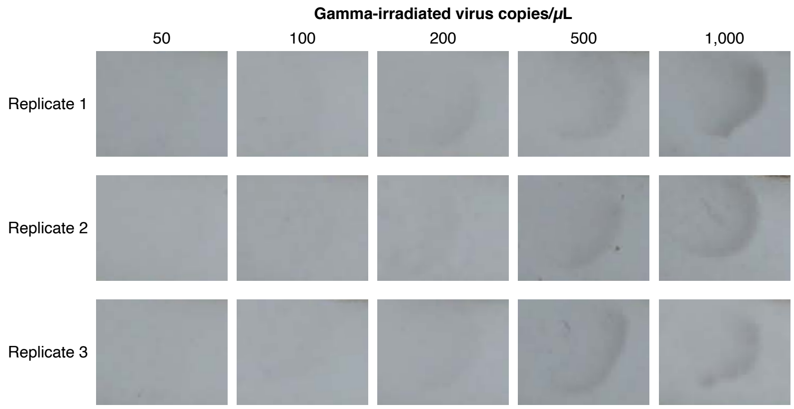


**Figure S16: Viral RNA detection: sensitivity of amplified HCR lateral flow assay (cf. Figure 4a).** Following the methods of Section S1.4.3, gamma-irradiated SARS-CoV-2 was spiked into extraction buffer with DNA probes and heated at 65 °C for 15 min; this solution was added to the Channel 1 well in a 96-well plate and the ends of the three nitrocellulose channels (Channel 1, Channel 2, Channel 3) were simultaneously submerged in their respective wells in the 96-well plate to start the test. The test region was photographed after 90 min. $N = 3$ replicate assays at each target concentration. The test region is visible in all three replicates down to a limit of detection of 200 copies/$\mu$L.

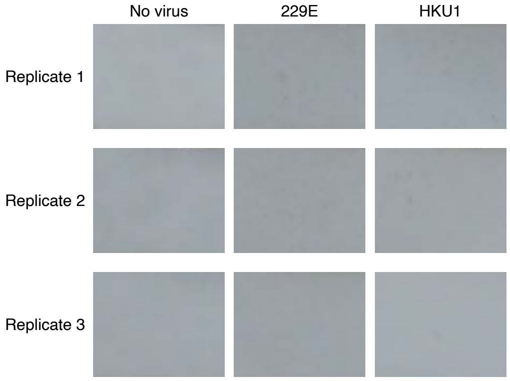


**Figure S17: Viral RNA detection: background and cross-reactivity of amplified HCR lateral flow assay (cf. Figure 4bc).** Following the methods of Section S1.4.3, no virus (to measure background) or off-target synthetic RNA genomes from coronaviruses 229E or HKU1 (to test cross-reactivity) were spiked into extraction buffer with DNA probes and heated at 65 °C for 15 min; this solution was added to the Channel 1 well in a 96-well plate and the ends of the three nitrocellulose channels (Channel 1, Channel 2, Channel 3) were simultaneously submerged in their respective wells in the 96-well plate to start the test. For the cross-reactivity tests, the off-target viral RNA genomes were spiked in at high concentration (7,200 copies/$\mu$L for 229E and 10,000 copies/$\mu$L for HKU1). The test region was photographed after 90 min. $N = 3$ replicate assays at each target concentration. For each target type, no staining was visible at the test region for all three replicates, indicating that there is no visible background when SARS-CoV-2 is absent, and no visible cross-reactivity with off-target 229E or HKU1 coronavirus RNA genomes.

### S3 Additional studies

#### S3.1 Gel study of HCR polymerization at short time scales

The gel study of Figure S18 demonstrates that an HCR initiator can trigger self-assembly of HCR polymers in excess of $\approx$20,000 bp ($>$ 500 HCR hairpins) in 10 minutes with HCR hairpins at 0.5 $\mu$M, suggesting the potential for achieving signal amplification of up to two orders of magnitude using HCR in the context of rapid lateral flow assays.

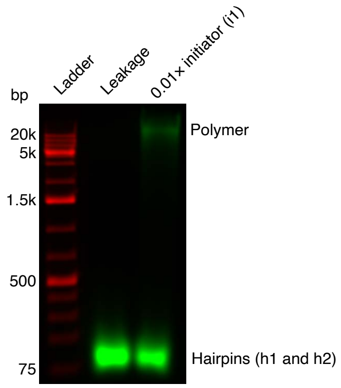


**Figure S18: Characterization of HCR polymerization at short time scales via agarose gel electrophoresis.** Following the methods of Section S1.2, HCR initiator (i1) triggers the self-assembly of HCR hairpins (h1 and h2) into amplification polymers. Each HCR hairpin (h1 and h2) at 0.5 $\mu$M with initiator i1 at 0.01$\times$. Polymerization time: 10 min. Initiator i1 is omitted from the leakage lane to demonstrate that HCR hairpins are kinetically trapped and do not polymerize in the absence of initiator. Green channel: fluorescence from Alexa647-labeled HCR hairpins h1 and h2 (displayed with 1% of pixels saturated). Red channel: GeneRuler 1 kb Plus DNA Ladder pre-stained with SYBR Gold.

#### S3.2 Measurement of HCR polymer length and amplification gain in the context of lateral flow assays

Following the methods of Section S1.5, here we characterize HCR polymer length and amplification gain in the context of lateral flow assays as follows:

- **HCR polymer length:** Using Alexa647-labeled HCR hairpins, we compare fluorescent signal using both h1 and h2 (permitting growth of HCR amplification polymers) vs with h1 only (permitting only a single h1 binding event with no polymerization due to the absence of h2). The ratio of the these two signal intensities provides a measurement of HCR polymer length in the context of the lateral flow assay format (in this case, for viral protein detection). The estimated HCR polymer length is $\approx 40$ (Table S1).
- **HCR amplification gain:** Using DIG-labeled HCR hairpins and CB-labeled anti-DIG reporter antibodies, we compare the CB signal using both h1 and h2 (permitting growth of HCR amplification polymers) vs with h1 only (permitting only a single h1 binding event with no polymerization due to the absence of h2). The ratio of the these two signal intensities provides a measurement of HCR amplification gain in the context of the lateral flow assay format (for either viral protein detection (Figure S20) or viral RNA detection (Figure S21)). The measured HCR amplification gain is $\approx$14 in the viral protein assay and $\approx$10 in the viral RNA assay (Table S2).

The fact that the amplification gain using CB-labeled anti-DIG reporter antibodies is lower than the HCR polymer length measured without using reporter antibodies suggests that there is room for improvement in optimizing the interaction between reporter antibodies and amplification polymers (e.g., to alleviate potential molecular crowding caused by bulky CB labels). Ideally, the amplification gain would match the mean polymer length, corresponding to a situation in which a reporter antibody is binding to each hairpin within an amplification polymer. With further optimization, it is plausible that the mean polymer length and amplification gain could be increased to two orders of magnitude within the constraints of the lateral flow assay format and a 1 hour overall assay duration.

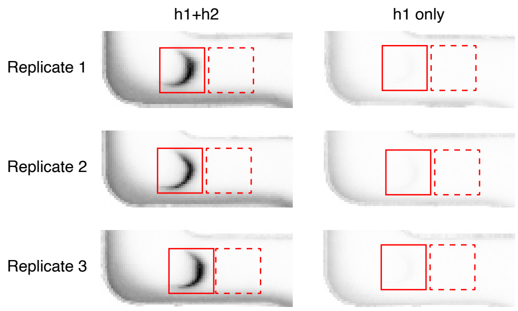


**Figure S19: Measurement of HCR polymer length in the context of a lateral flow assay for viral protein detection.** Following the methods of Section S1.5.1, two types of experiment are compared: amplified assays (using both h1 and h2, so that HCR polymerization can proceed) vs unamplified assays (using h1 only, permitting only a single h1 binding event with no polymerization due to the absence of h2). Gamma-irradiated virus was spiked into a mixture of saliva and extraction buffer at 20,000 copies/$\mu$L for all tests. $N = 3$ replicate assays for each condition. Quantitative image analysis following the methods of Section S1.5.3 using the depicted signal boxes (solid boundary) and background boxes (dashed boundary).

| | $\text{Signal}_{\text{h1+h2}}$ | $\text{Signal}_{\text{h1}}$ | Polymer length |
|---|---|---|---|
| Viral protein detection assay | $1070 \pm 16$ | $26 \pm 5$ | $41 \pm 8$ |

**Table S1: Estimated HCR polymer length in the context of a lateral flow assay for viral protein detection.** Quantitative image analysis of the amplified (h1 and h2) and unamplified (h1 only) assays of Figure S19 following the methods of Section S1.5.3. Mean $\pm$ estimated standard error of the mean via uncertainty propagation for $N = 3$ replicate assays for each experiment type.

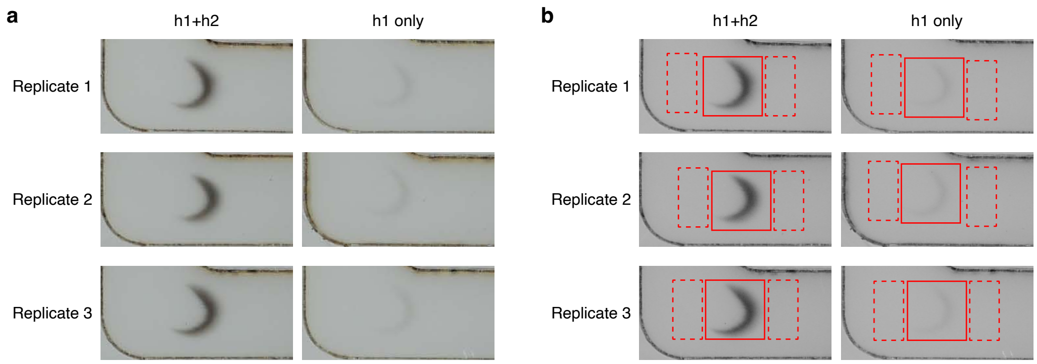


**Figure S20: Measurement of HCR amplification gain in the context of a lateral flow assay for viral protein detection.** Following the methods of Section S1.5.2, two types of experiment are compared: amplified assays (using both h1 and h2, so that HCR polymerization can proceed) vs unamplified assays (using h1 only, permitting only a single h1 binding event with no polymerization due to the absence of h2). Gamma-irradiated virus was spiked into a mixture of saliva and extraction buffer at 5000 copies/$\mu$L for all tests. $N = 3$ replicate assays for each condition. (a) Raw images. (b) Images after conversion to grayscale. Quantitative image analysis following the methods of Section S1.5.3 using the depicted signal boxes (solid boundary) and background boxes (dashed boundary).

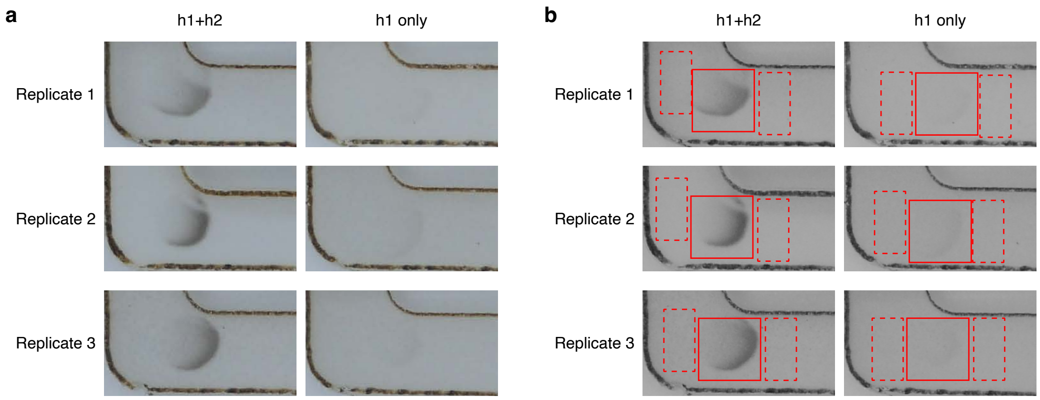


**Figure S21: Measurement of HCR amplification gain in the context of a lateral flow assay for viral RNA detection.** Following the methods of Section S1.5.2, two types of experiment are compared: amplified assays (using both h1 and h2, so that HCR polymerization can proceed) vs unamplified assays (using h1 only, permitting only a single h1 binding event with no polymerization due to the absence of h2). Gamma-irradiated virus was spiked into extraction buffer at 5000 copies/$\mu$L for all tests. $N = 3$ replicate assays for each condition. (a) Raw images. (b) Images after conversion to grayscale. Quantitative image analysis following the methods of Section S1.5.3 using the depicted signal boxes (solid boundary) and background boxes (dashed boundary).

| | $\text{Signal}_{\text{h1+h2}}$ | $\text{Signal}_{\text{h1}}$ | Amplification gain |
|---|---|---|---|
| Viral protein detection assay | $19.6 \pm 0.8$ | $1.43 \pm 0.05$ | $13.7 \pm 0.8$ |
| Viral RNA detection assay | $13 \pm 2$ | $1.3 \pm 0.3$ | $10 \pm 3$ |

**Table S2: Estimated amplification gain in the context of lateral flow assays for viral protein or RNA detection.** Quantitative image analysis of the amplified (h1 and h2) and unamplified (h1 only) assays of Figures S20 and S21 following the methods of Section S1.5.3. Mean $\pm$ estimated standard error of the mean via uncertainty propagation for $N = 3$ replicate assays for each experiment type.

#### S3.3 Visualization of automated reagent delivery using a 3-channel lateral flow assay with food coloring

Food coloring was used to validate that the 3-channel nitrocellulose membranes for the viral protein assay and viral RNA assay lead to successive delivery from Channels 1, 2, and 3 to the test region.

- **Viral protein detection:** For the protein detection assay format, green, blue, and red food coloring was diluted 1:30 in 5× SSC and 0.1% Tween-20. Diluted food coloring was added to unblocked conjugate pads. Diluted green food coloring (75 $\mu$L) was added to the Channel 1 conjugate pad, diluted blue food coloring (50 $\mu$L) was added to the Channel 2 conjugate pad, and diluted red food coloring (100 $\mu$L) was added to the Channel 3 conjugate pad. The conjugate pads were dried in a 37 °C oven for 1 h before mounting to the folding card device. Lastly, 300 $\mu$L 5× SSC with 0.1% Tween-20 was added to the sample pad before closing the folding card device, initiating the flow of liquid from the three conjugate pads onto the nitrocellulose membrane (Supplementary Video 1).
- **Viral RNA detection:** For the RNA detection assay format, 100 $\mu$L green, blue, and red food coloring (diluted 1:200 in 5× SSC and 0.1% Tween-20) was added to the Channel 1, 2, and 3 wells, respectively, and the ends of the three nitrocellulose channels were simultaneously submerged in their respective wells to initiate the flow of liquid to the test region (Supplementary Video 2).

For both assay formats, successive flow of the three colors demonstrates that liquid from each of the three channels reaches the test region in the correct order.
